# Selective degradation of multimeric proteins via chemically induced proximity to TRIM21

**DOI:** 10.1101/2024.01.31.578122

**Authors:** Panrui Lu, Yalong Cheng, Lei Xue, Xintong Ren, Chenglong Chen, Jiao Li, Qingcui Wu, Shan Sun, Junjie Hou, Wei Jia, Chao Li, Xiangbing Qi, Niu Huang, Ting Han

**Affiliations:** Tsinghua Institute of Multidisciplinary Biomedical Research, Tsinghua University, Beijing, 102206, China; National Institute of Biological Sciences, Beijing, 102206, China; College of Life Sciences, Beijing Normal University, Beijing, 100875, China; State Key Laboratory of Membrane Biology, Beijing Frontier Research Center for Biological Structures, School of Life Sciences, Tsinghua University, Beijing, 100084, China; Deepkinase Co, Ltd, Beijing, 102206, China

## Abstract

Targeted protein degradation (TPD) has emerged as an effective strategy to eliminate disease-causing proteins by inducing their interactions with the protein degradation machinery. First-generation TPD agents exploit a limited set of broadly expressed E3 ubiquitin ligases with constitutive activity, forbidding their application to proteins requiring higher levels of targeting selectivity. Here, by phenotype-based screening, we discovered that the antipsychotic drug acepromazine possesses interferon-enhanced cytotoxicity towards cancer cell lines expressing high levels of aldo-keto reductases 1C. These enzymes convert acepromazine into its stereo-selective metabolite (*S*)-hydroxyl-acepromazine, which recruits the interferon-induced E3 ubiquitin ligase TRIM21 to the vicinity of the nuclear pore complex, resulting in the degradation of nuclear pore proteins. Co-crystal structures of acepromazine and derivatives in complex with the PRYSPRY domain of TRIM21 revealed a ligandable pocket, which was exploited for designing heterobifunctional degraders. The resulting chemicals selectively degrade multimeric proteins— such as those in biomolecular condensates—without affecting monomeric proteins, consistent with the requirement of substrate-induced clustering for TRIM21 activation. As aberrant protein assemblies have been causally linked to diseases such as neurodegeneration, autoimmunity, and cancer, our findings highlight the potential of TRIM21-based multimer-selective degraders as a strategy to tackle the direct causes of these diseases.

## Introduction

Targeted protein degradation (TPD) via chemically induced proximity between E3 ubiquitin ligases and neo-substrate proteins is an emerging therapeutic modality for tackling proteins deemed undruggable with traditional small molecule inhibitors^1^. Two categories of chemicals have been leveraged for TPD, including monovalent molecular glue degraders and bifunctional PROTACs (proteolysis targeting chimeras)^2,3^. The impact of TPD is highlighted by IMiDs (immunomodulatory imide drugs), which function as molecular glues to induce the degradation of multiple neo-substrates, leading to exceptional efficacy in treating multiple myeloma^4,5^. Recently, a plethora of rationally designed PROTACs targeting disease-causing proteins have entered clinical trials and provided proof-of-concept for the efficacy and safety of TPD in human patients^3^. However, current TPD strategies mainly exploit E3 ligase complexes containing cereblon (CRBN) or von Hippel−Lindau (VHL), despite the presence of over 600 E3 ligases in human cells^6^. Thus, chemicals that employ additional E3 ligases with differentiated properties are highly sought after to avoid the limitations of current TPD agents and broaden the scope of TPD applications to unmet medical needs.

First-generation TPD agents utilize E3 ligases with general expression patterns and constitutive activity, and are thus unsuitable for targeting proteins with essential functions in healthy tissues^7^. Several strategies are actively being explored to increase TPD selectivity. For example, the nuclear localization of the E3 ubiquitin ligase DCAF16 has been harnessed to restrict TPD to the nucleus^8^. Moreover, E3 ubiquitin ligases with tissue-selective expression are enthusiastically pursued to restrict TPD to specific diseased tissues^9,10^. Many E3 ligases are regulated by autoinhibitory mechanisms either by hindering the recruitment of E2-ubiquitin complexes or constraining the structural flexibility of the E3 ligases^11^. Such mechanisms enhance the specificity of these E3 ligases towards their cognate substrates, but have so far not been exploited to improve TPD selectivity.

One area where selective TPD would be an attractive technology is the therapeutic targeting of biomolecular condensates, which are membraneless assemblies enriching specific proteins and nucleic acids in subcellular environments^12^. An explosion of recent studies unveils key functions of biomolecular condensates in normal and diseased states^13^. In particular, aberrant protein condensation has been causally linked to neurodegeneration, autoimmunity, and cancer, thus revealing exciting therapeutic opportunities^13^. However, selective degradation of proteins in biomolecular condensates without affecting those in the dilute phase has proven to be technically challenging^14^.

Here, we present TRIM21-mediated TPD as a general strategy to induce selective degradation of multimeric proteins, including those in biomolecular condensates. By phenotype-based screening followed by target deconvolution, we discovered (*S*)-hydroxyl-acepromazine as a monovalent degrader of several proteins in the multimeric nuclear pore complex via the E3 ubiquitin ligase TRIM21. Co-crystal structures of the PRYSPRY domain of TRIM21 in complex with acepromazine and its metabolites revealed a ligandable pocket, which was exploited for PROTAC design. The resulting TRIM21-based PROTACs are highly selective degraders for multimeric targets in biomolecular condensates without degrading non-multimeric targets in the dilute phase. These findings highlight the potential of TRIM21-based degraders to eliminate target proteins that cause disease via aberrant multimerization.

## Results

### Acepromazine displays interferon-enhanced selective anticancer activity

We sought to identify small molecules with enhanced cytotoxicity to cancer cells in the presence of IFNγ, which is a key inflammatory cytokine in the tumor microenvironment that orchestrates diverse antitumor responses^15^. Using the lung adenocarcinoma cell line A549, we designed an in vitro competitive cell growth assay: first, we used CRISPR/Cas9 to delete *IFNGR1* (encoding a subunit of the IFNγ receptor) from A549 cells, resulting in a pool of cells unable to sense IFNγ (Figure S1A); second, we expressed histone H2B-GFP and H2B-mCherry fusion proteins in wild-type (WT) and *IFNGR1*-deficient A549 cells, respectively; third, we mixed the two engineered cell populations at a 1:1 ratio, treated them with IFNγ and small molecules from an in-house chemical library for four days, and then monitored the changes in the ratio of green to red nuclei (Figure 1A).

**Figure 1:**
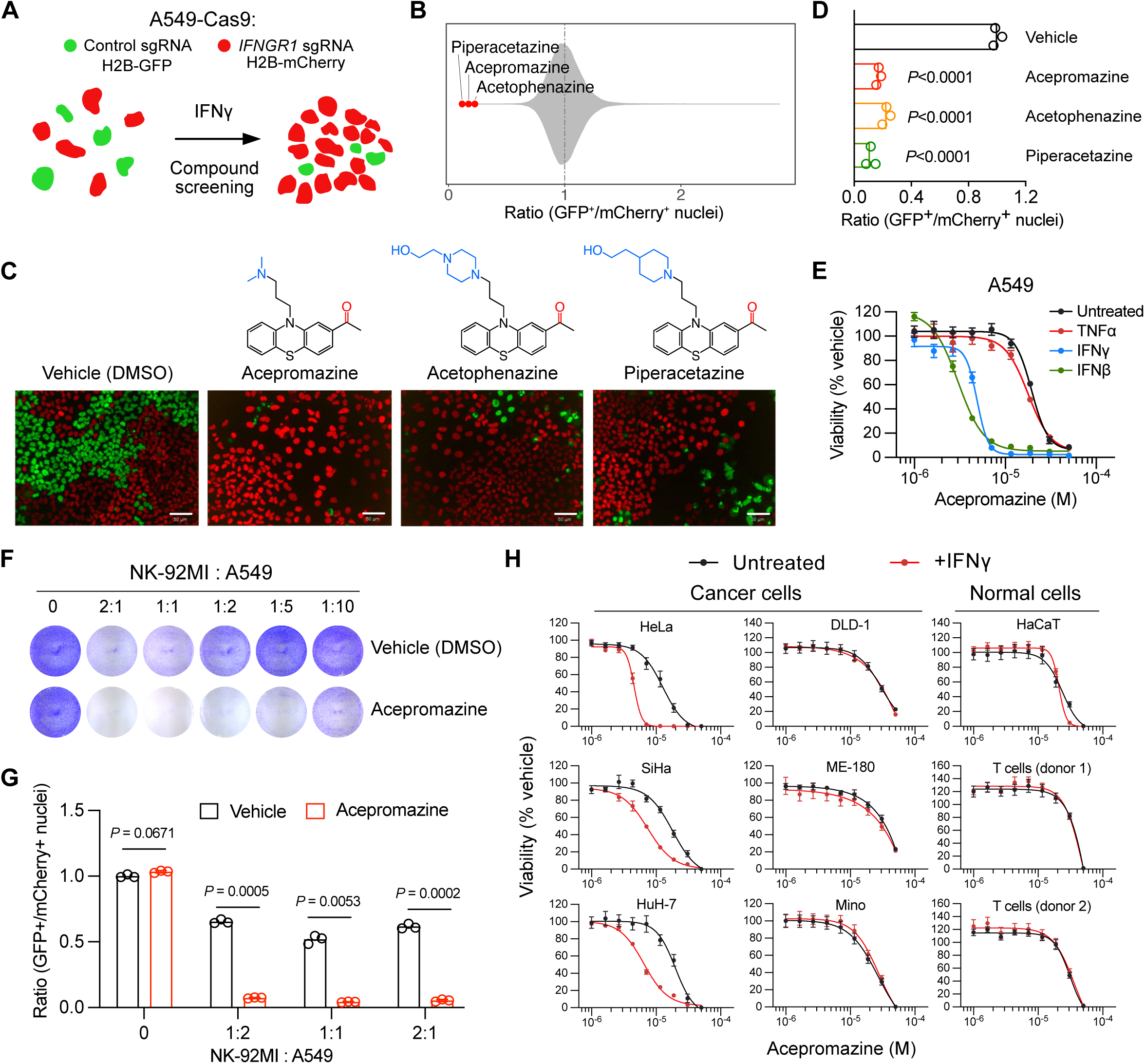
Acepromazine displays an interferon-enhanced selective anticancer activity. (A) Schematic of the competitive cell growth assay. An equal mixture of WT A549 cells (labeled with H2B-GFP) and *IFNGR1*-deficient A549 cells (labeled with H2B-mCherry) were treated with IFNγ (20 ng/ml) and test compounds (10 µM) for four days followed by high-content imaging to quantify the ratio of GFP^+^ nuclei to mCherry^+^ nuclei. (B) Effects of 81,845 compounds on the ratio of GFP^+^ nuclei to mCherry^+^ nuclei in the primary screen. (C) Chemical structures of hit compounds and their representative raw images from the primary screen. Green: H2B-GFP; Red: H2B-mCherry; Scale bar: 50 µm. (D) Effects of hit compounds on the ratio of GFP^+^ nuclei to mCherry^+^ nuclei. Data represent the mean of three independent samples. One-way ANOVA followed by Dunnett’s multiple comparison test was used to determine statistical significance. (E) Concentration-response curves of acepromazine on the viability of A549 cells treated with indicated cytokines (10 ng/ml). Data represent the mean ± s.e.m. of three independent samples. IC_50_ and 95% CI are shown in Table S1. (F) Crystal violet staining of A549 cells co-cultured with indicated amounts of NK-92MI cells with or without acepromazine (10 µM) treatment for 48 h. A representative result was shown from two independent experiments. (G) Effects of NK-92MI cells on the fitness of WT A549 cells (labeled with H2B-GFP) relative to *IFNGR1*-deficient A549 cells (labeled with H2B-mCherry) with or without acepromazine (10 µM) treatment for 48 h. The ratios of GFP^+^ nuclei to mCherry^+^ nuclei were determined by FACS. Data represent the mean of three independent samples. Unpaired Student’s t-tests (two-tailed) were used to determine statistical significance. (H) Concentration-response curves of acepromazine on the viability of indicated cell lines and primary T lymphocytes with or without IFNγ (10 ng/ml) treatment. Data represent the mean ± s.e.m. of three independent samples. IC_50_ and 95% CI are shown in Table S1.

By screening 81,845 small drug-like molecules (at 10 µM), we identified three hits— acepromazine, acetophenazine, and piperacetazine—that selectively reduced the fitness of WT A549 cells relative to *IFNGR1*-deficient A549 cells (indicated by reduced ratios of green to red nuclei) (Figures 1B-1D). The three hit molecules share a common acetyl-substituted phenothiazine core linked to different amine-containing moieties by a propyl chain (Figure 1C). Due to their structural similarity, we focused on acepromazine, hereafter referred to as ACE, for further characterization. By measuring A549 viability, we observed the enhancement of ACE cytotoxicity by both IFNγ and interferon beta (IFNβ), but not by tumor necrosis factor alpha (TNFα) (Figure 1E). To mimic the sources of IFNγ production in the tumor microenvironment, we utilized the natural killer (NK) cell line NK-92MI, which secretes IFNγ upon encountering with cancer cells^16^. We found that ACE cytotoxicity against A549 cells could be enhanced by coculturing with NK-92MI. Such enhancement was lost in *IFNGR1*-deficient A549 cells (Figures 1F-1G and S1B).

We further examined the cytotoxicity of ACE across six cancer cell lines, one normal cell line, and activated T lymphocytes from two healthy donors. IFNγ-enhanced cytotoxicity was observed in the liver cancer cell line HuH-7 and the cervical cancer cell lines HeLa and SiHa. In contrast, no IFNγ-enhanced cytotoxicity was observed in the lymphoma cell line Mino, the colorectal cancer cell line DLD-1, and the cervical cancer cell line ME-180. Moreover, the non-cancerous keratinocyte cell line HaCat and activated T lymphocytes from healthy donors were resistant to IFNγ-enhanced ACE cytotoxicity. These data, taken together, reveal that ACE possesses an interferon-enhanced anticancer activity towards a subset of cancer cell lines.

### Metabolic activation of acepromazine underlies its selective anticancer activity

ACE is an antipsychotic drug used in humans during the 1950s, but is currently used only as a sedative in veterinary medicine^17^. The behavioral effect of ACE is primarily attributed to antagonism of post-synaptic dopamine receptors^18^. However, chlorpromazine (Figure S1C), a closely related analog that is still used as an antipsychotic drug in humans^19^, did not display interferon-enhanced anticancer activity (Figure S1D). Moreover, ACE-sensitive cancer cell lines do not express dopamine receptors (Figure S1E). These findings indicate that ACE exerts its anticancer activity independent of dopamine receptors.

To elucidate the mechanism of action of ACE, we first focused on its selectivity towards a subset of cancer cell lines. As ACE is known to be susceptible to metabolic conversions^20^, we hypothesized that ACE might be an inactive pro-drug activated by a subset of cancer cell lines. To test this hypothesis, we treated the sensitive cell line A549 with ACE and then used the conditioned medium to treat the insensitive cell line DLD-1 (Figure 2A). While untreated ACE was not toxic to DLD-1, ACE following A549 conditioning exhibited potent cytotoxicity. Heating did not inactive the cytotoxic activity in the conditioned medium, suggesting that the active ingredient is a small-molecule metabolite (Figure 2B). We thus extracted total metabolites from conditioned medium of both ACE-sensitive and insensitive cell lines and resolved them on thin-layer chromatography (TLC). Ultraviolet-light illumination of TLC revealed time-dependent conversion of ACE into a more polar species in sensitive cell lines (A549 and HeLa), but not in insensitive cell lines (DLD-1 and HCT-116) (Figure 2C). As the carbonyl group of ACE is known to be susceptible to reduction, we chemically reduced ACE to ACE-OH (hydroxyethylpromazine) and used it as a reference. The TLC pattern of the more polar species matched ACE-OH (Figure 2C). Moreover, liquid chromatography–mass spectrometry analyses confirmed the new species differed from ACE by 2 Daltons, corresponding to the addition of two hydrogen atoms (Figures S2A-S2B). These results revealed selective conversion of ACE into ACE-OH by sensitive cancer cell lines.

**Figure 2:**
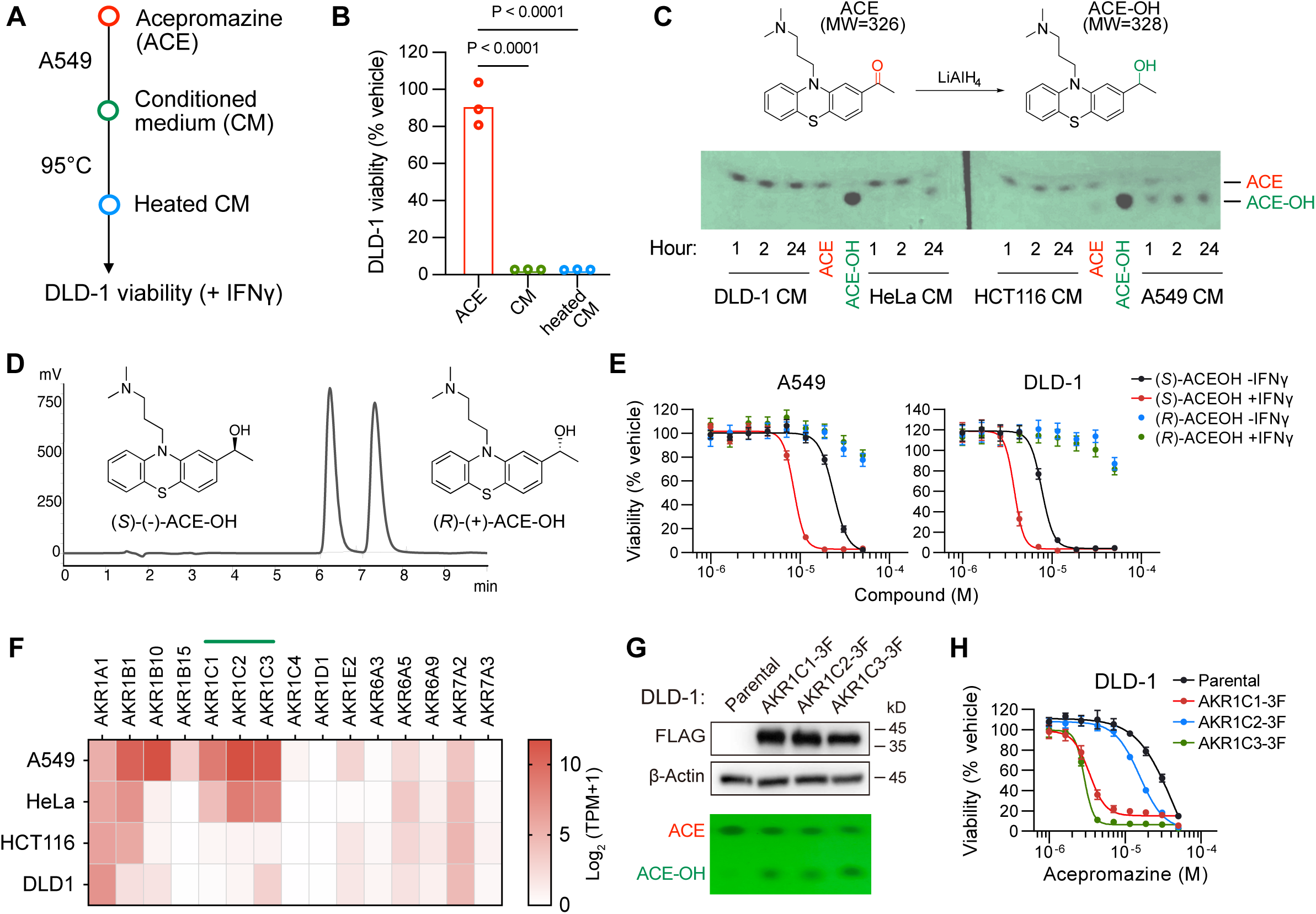
Metabolic activation of acepromazine underlies its selective anticancer activity. (A) Workflow for the preparation of conditioned medium (CM). (B) The effects of ACE, CM, and heated CM on the viability of DLD-1 (treated with 10 ng/ml IFNγ) for 3 days. Data represent the mean of three independent samples. One-way ANOVA followed by Dunnett’s multiple comparison test was used to determine statistical significance. (C) Chemical reduction of ACE to ACE-OH. (D) Detection of time-dependent conversion of ACE to ACE-OH in indicated cell lines by TLC. (E) Separation of indicated ACE-OH enantiomers by chiral column chromatography. (F) Concentration-response curves of (*S*)-ACE-OH and (*R*)-ACE-OH on the viability of indicated cell lines with or without IFNγ (10 ng/ml) treatment. Data represent the mean ± s.e.m. of three independent samples. IC_50_ and 95% CI are shown in Table S1. (G) Heatmap showing the expression levels of genes encoding aldo-keto reductases in indicated cell lines. TPM (transcripts per million) values were downloaded from DepMap. (H) Detection of ectopically expressed AKR1C1/2/3-3xFLAG in DLD-1 cells by anti-FLAG western blotting and ACE to ACE-OH conversion by TLC. A representative result was shown from three independent experiments. Uncropped western blot images are provided as a Source Data file. (I) Concentration-response curves of ACE on the viability of DLD-1 cells (treated with 10 ng/ml IFNγ) expressing AKR1C1/2/3-3xFLAG. Data represent the mean ± s.e.m. of three independent samples. IC_50_ and 95% CI are shown in Table S1.

The conversion of ACE into ACE-OH created a chiral center. As enantiomers of chiral drugs can have reduced, no, or better effects than racemates, we separated the ACE-OH racemate into two enantiomer fractions by chiral column chromatography (Figure 2D). By chemical derivation to facilitate crystallization followed by X-ray diffraction, we determined the absolute conformations and optical activities of the two enantiomers to be (*S*)-(-)-ACE-OH and (*R*)-(+)-ACE-OH (Figures S2C-S2D). We then treated A549 and DLD-1 cells with the purified ACE-OH enantiomers. (*S*)-ACE-OH displayed IFNγ-enhanced toxicity to both cell lines. In contrast, (*R*)-ACE-OH was inactive in either cell line (Figure 2E). Thus, we conclude that (*S*)-ACE-OH is the active metabolite of ACE.

The aldo-keto reductase (AKR) family of enzymes catalyze the reduction of the carbonyl groups of various endogenous and exogenous substrates^21^. We therefore compared the expression levels of genes encoding aldo-keto reductases in ACE-converting versus non-converting cell lines and observed that high levels of AKR1C1, AKR1C2, and AKR1C3 were associated with ACE to ACE-OH conversion (Figure 2F). We then individually expressed AKR1C1, AKR1C2, and AKR1C3 in DLD-1 cells and found they all enabled DLD-1 to convert ACE into ACE-OH (Figure 2G). Moreover, expression of AKR1C1, AKR1C2, or AKR1C3 rendered DLD-1 cell more sensitive to ACE (Figure 2H). Similar results were obtained in another ACE-insensitive cell line ME-180 (Figures S2E-S2F). Notably, AKR1C1/2/3 are overexpressed in cholangiocarcinoma, liver hepatocellular carcinoma, and lung squamous cell carcinoma (Figure S2G). These observations not only explain the observed anticancer selectivity of ACE, but also suggest that activation of ACE by tumor-associated AKR1C1/2/3 could be a strategy to enrich cytotoxic metabolites in tumors.

### Interferon-induced TRIM21 mediates the anticancer activity of acepromazine

Next, we sought to investigate the mechanism by which interferons enhance ACE’s anticancer activity. We performed pooled genome-wide CRISPR-Cas9 knockout screening by targeting 19,114 genes with four individual sgRNAs per gene^22^. We treated sgRNA-transduced A549 cells with either IFNγ or IFNγ plus a sub-lethal dose of ACE for three weeks. Afterwards, we isolated genomic DNAs from surviving cells and performed next-generation sequencing to measure the abundance of each sgRNA. Using the MAGeCK algorithm^23^, we found six top-ranking enriched genes, included five involved in IFNγ sensing and signal transduction (*IFNGR1*, *IFNGR2*, *JAK1*, *JAK2*, and *STAT1*) as well as *TRIM21* (Figures 3A-3B).

**Figure 3:**
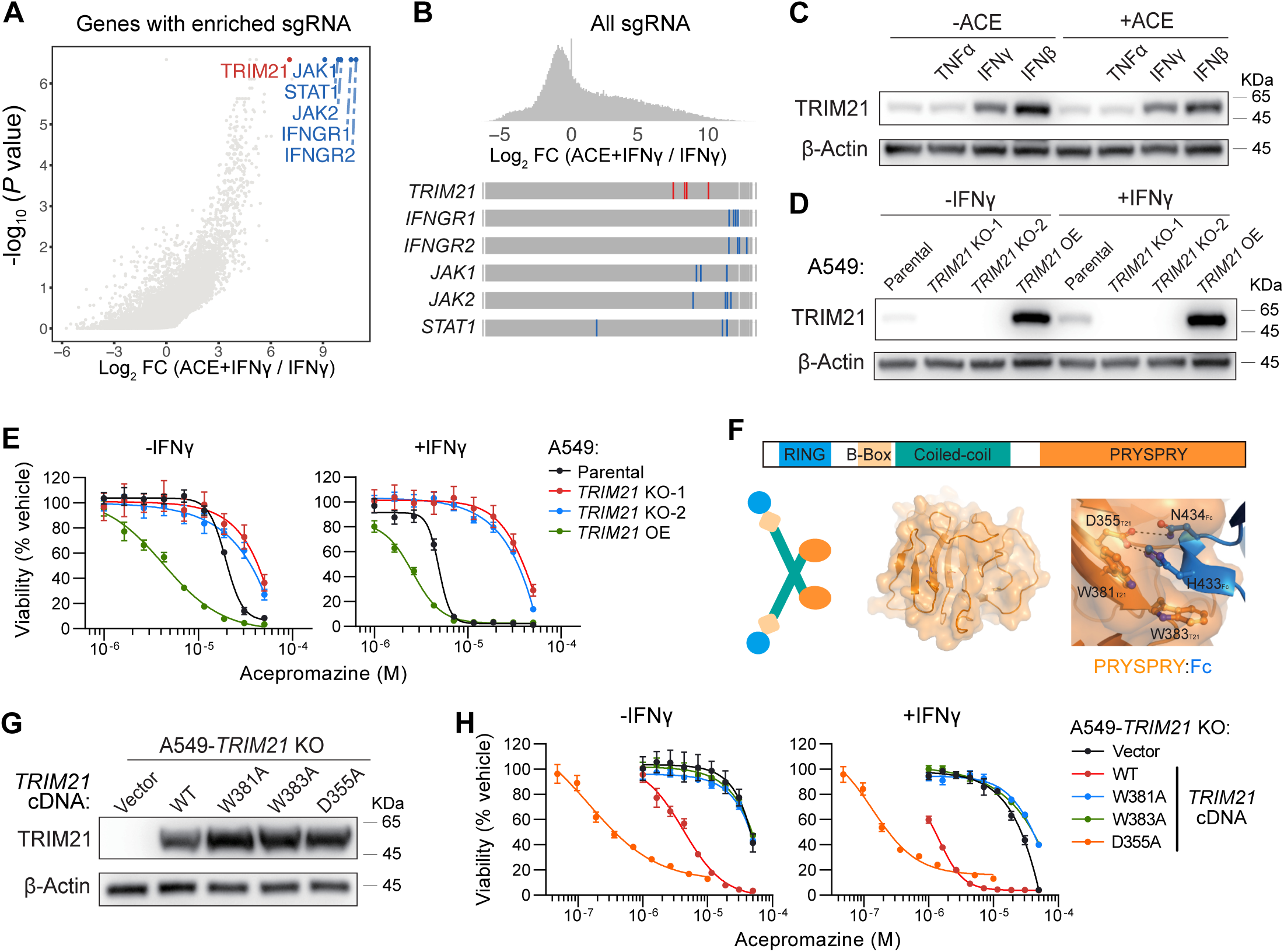
Interferon-induced TRIM21 mediates the anticancer activity of acepromazine. (A) Scatterplot depicting genes with significantly enriched sgRNAs in A549-Cas9 cells treated with ACE (4-10 µM) and IFNγ (10 ng/ml) versus IFNγ (10 ng/ml) only (n = 3 per group) for 3 weeks. *P* values were computed by MAGeCK from the negative binomial model using a modified robust ranking aggregation algorithm. Data used for generating the plot are provided in Table S2. (B) Frequency histogram of all sgRNAs and ranks of sgRNAs targeting indicated genes. (C) Immunoblots of TRIM21 and β-Actin in A549 cells pre-treated with indicated cytokines (10 ng/ml) with or without ACE (10 µM) treatment for 12 h. A representative result was shown from three independent experiments. Uncropped western blot images are provided as a Source Data file. (D) Immunoblots of TRIM21 and β-Actin in A549 cells of indicated genotypes with or without IFNγ (10 ng/ml) treatment for 12 h. A representative result was shown from three independent experiments. Uncropped western blot images are provided as a Source Data file. (E) Concentration-response curves of ACE on the viability of A549 cells with indicated genotypes with or without IFNγ (10 ng/ml) treatment. Data represent the mean ± s.e.m. of three independent samples. IC50 and 95% CI are shown in Table S1. (F) Domain architecture of TRIM21. (G) Crystal structure of the PRYSPRY domain of TRIM21 in complex with Fc (modified from PDB: 2IWG). (H) Immunoblots of TRIM21 and β-Actin in A549-*TRIM21* knockout (KO) cells rescued with indicated *TRIM21* cDNAs. A representative result was shown from three independent experiments. Uncropped western blot images are provided as a Source Data file. (I) Concentration-response curves of ACE on the viability of A549-*TRIM21* KO cells rescued with indicated *TRIM21* cDNAs with or without IFNγ (10 ng/ml) treatment. Data represent the mean ± s.e.m. of three independent samples. IC50 and 95% CI are shown in Table S1.

*TRIM21* encodes a tripartite motif-containing E3 ubiquitin ligase, whose expression is induced by interferons^24,25^. In line with previous findings, western blotting of A549 lysates confirmed that TRIM21 could be induced by IFNβ or IFNγ, but not by TNFα (Figure 3C). We thus hypothesized that interferons enhance ACE’s anticancer activity by inducing the expression of TRIM21. To test this hypothesis, we generated two A549 *TRIM21* knockout clones. In addition, we overexpressed TRIM21 in A549 cells to mimic interferon-induced expression of TRIM21 (Figure 3D). By measuring cell viability, we found that loss of TRIM21 rendered A549 cells resistant to ACE, whereas TRIM21 overexpression resulted in enhanced sensitivity to ACE, thus overriding the effect of IFNγ (Figure 3E). Similar results were obtained in two additional ACE-sensitive cell lines SiHa and HuH-7 (Figures S3A-S3B). Based on these results, we conclude that interferon-induced TRIM21 mediates the anticancer activity of ACE.

TRIM21 is an innate immune sensor, which recognizes cytosolic antibody-coated viruses to trigger antiviral responses^26,27^. The C-terminal PRYSPRY domain of TRIM21 has a hydrophobic pocket that binds the fragment crystallizable (Fc) region of antibodies^28^ (Figure 3F). We selected three residues in this pocket important for Fc binding and expressed their alanine substitution mutants (D355A, W381A, and W383A) in *TRIM21* knockout cells (Figure 3G). The W381A and W383A mutants could not restore ACE sensitivity. In contrast, the D355A mutant displayed increased ACE sensitivity in comparison to WT TRIM21 (Figure 3H). The identification of both loss- and gain-of-activity mutations suggests that this pocket of TRIM21^PRYSPRY^ is involved in the protein-compound or protein-protein interactions that mediate ACE activity.

### Acepromazine induces TRIM21-dependent degradation of nuclear pore proteins

As TRIM21 possesses an N-terminal RING domain with E3 ubiquitin ligase activity, we hypothesized that ACE functions by directing TRIM21 to degrade some proteins required for cell viability. To identify such proteins, we used label-free quantitative proteomics to compare the proteomes of A549 cells overexpressing TRIM21(D355A) treated with vehicle versus ACE (Figure 4A). We discovered several nuclear pore proteins—NUP35, NUP155, SMPD4, and GLE1—to be significantly depleted following ACE treatment. By western blotting, we found that ACE could trigger the degradation of these nuclear pore proteins in ACE-sensitive cell lines, but not in cell lines that are ACE-insensitive or *TRIM21*-deficient (Figures S4A-S4B). Moreover, inhibition of the proteasome (by bortezomib treatment) or the E1 ubiquitin activating enzyme (by TAK243 treatment) prevented ACE-induced degradation of these proteins (Figures 4B and S4C). As only the metabolite (*S*)-ACE-OH has anticancer activity, we examined whether ACE-OH could induce the degradation of nuclear pore proteins in a stereo-selective manner. Indeed, (*S*)-ACE-OH but not (*R*)-ACE-OH triggered the degradation of NUP35, NUP155, SMPD4, and GLE1 in IFNγ- stimulated A549 cells (Figure 4C).

**Figure 4:**
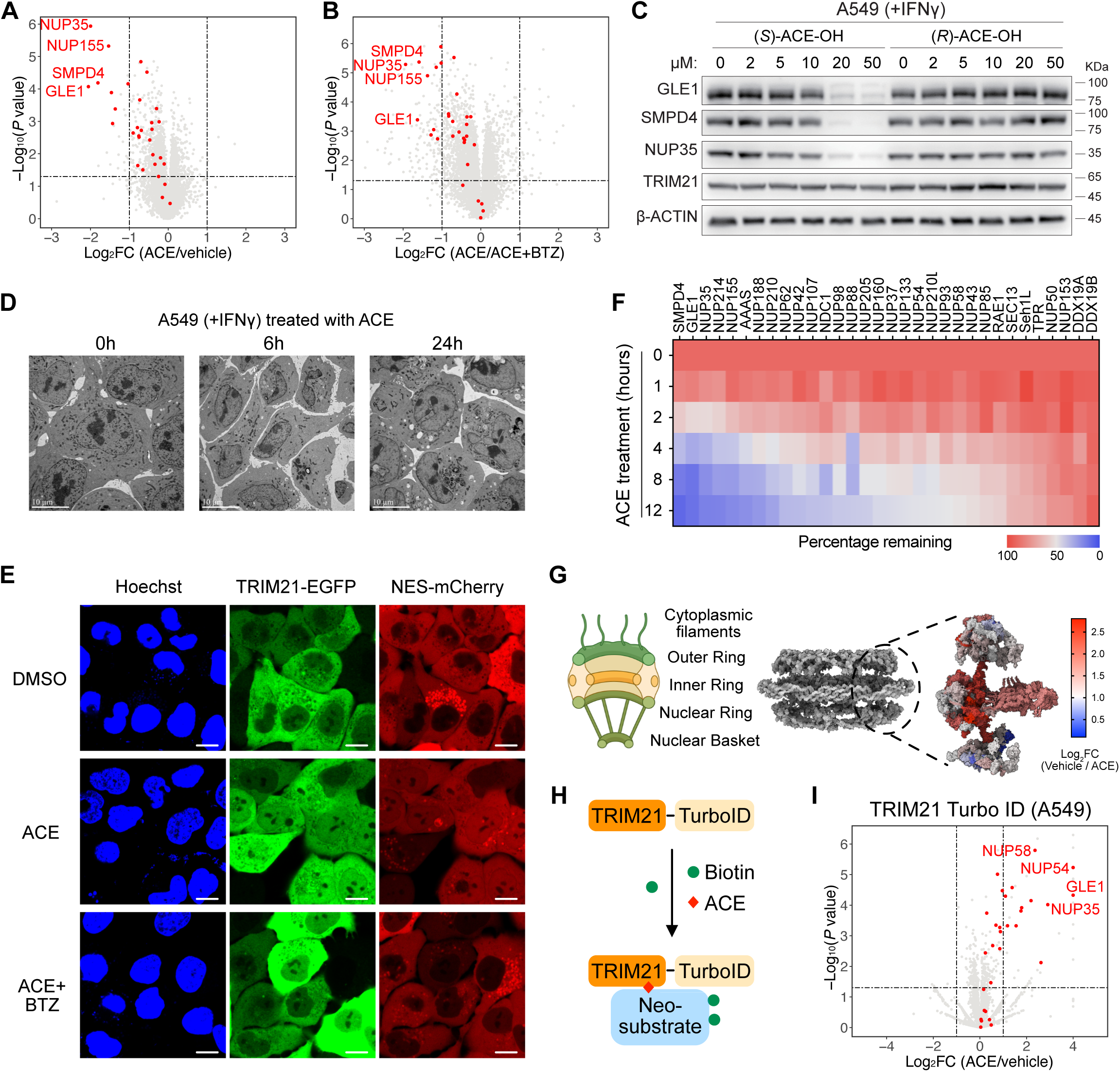
Acepromazine induces TRIM21-dependent degradation of nuclear pore proteins. (A-B) Volcano plots depicting log_2_ transformed average fold change of each protein (quantified by label-free proteomics) and −log_10_ transformed *P* value comparing A549-TRIM21^D355A^ treated with (A) ACE (10 µM) versus vehicle; (B) ACE (10 µM) versus ACE (10 µM)/BTZ (100 nM) for 8 h. BTZ: bortezomib. Three independent samples were included in the analyses. *P* values were calculated by unpaired Student’s t-test (two tailed). Data used for the plots are provided in Table S3. (C) Immunoblots of indicated proteins in IFNγ-pretreated A549 cells treated with (*S*)-ACE-OH or (*R*)-ACE-OH with or without for 8 h. A representative result was shown from three independent experiments. Uncropped western blot images are provided as a Source Data file. (D) Transmission electron microscopy images of IFNγ-pretreated A549 cells treated with ACE (10 µM) for indicated time. A representative result was shown from three independent experiments. Scale bar: 10 µm. (E) Confocal microscopy images of nuclei (blue, stained with Hoechst 33342), TRIM21-EGFP (green) and NES-mCherry (red) in A549 cells. When indicated, cells were pre-treated with BTZ (100 nM) for 2h followed by ACE (10 µM) treatment for 4h. A representative result was shown from three independent experiments. Scale bar: 10 µm. (F) Heatmap depicting time-dependent depletion of nuclear pore proteins (quantified by label-free proteomics) in IFNγ-stimulated A549 treated with ACE (10 µM). Data used for the plot are provided in Table S4. (G) Schematic cross-section of the NPC (Created with Biorender.com) and the degree of nuclear pore protein depletion mapped to the cryo-electron tomography structure of the NPC (PDB: 7R5J). (H) Schematic of the TRIM21-TurboID assay. (I) Volcano plot depicting log_2_ transformed average fold change of each protein (quantified by label-free proteomics) and −log_10_ transformed *P* value comparing TRIM21-TurboID-enriched proteins from A549 cells treated with ACE (20 µM) versus vehicle for 4 h. Three independent samples were included in the analyses. *P* values were calculated by unpaired Student’s t-test (two tailed). Data used for the plot are provided in Table S5.

The nuclear pore complex (NPC) is composed of ∼1000 building blocks assembled from copies of ∼30 proteins and organized as three stacked rings (the inner, cytoplasmic, and nuclear rings) flanked by the cytoplasmic filament and the nuclear basket^29^ (Figure 4D). By monitoring the abundance of each nuclear pore protein in ACE-treated A549 cells over time, we found that the inner ring of NPC was depleted rapidly (as short as four hours post ACE treatment), followed by the depletion of subunits from the cytoplasmic ring. Subunits specific to the nuclear ring and nuclear basket was not depleted over the 12-hour window (Figures 4D-4E). Consistent with the degradation of NPC subunits, high-resolution imaging by transmission electron microscopy revealed the distortion and fragmentation of nuclei in ACE-treated A549 cells (Figure 4F). Moreover, in A549 cells expressing GFP-tagged TRIM21 and NES (nuclear export signal)-tagged mCherry, ACE treatment resulted in the mislocalization of both proteins into the nucleus, consistent with the impairment of NPC function. ACE-dependent mis-localization of TRIM21-mEGFP and NES-mCherry could be rescued by proteasome inhibition, which is consistent with the involvement of protein degradation (Figure 4G). Taken together, these results suggest that ACE recruits TRIM21 from the cytosol to attack the inner ring of NPC, resulting in the functional impairment of nuclear pores.

To test the model of ACE-induced recruitment of TRIM21 to the NPC, we employed TRIM21 fused with the biotin ligase TurboID, which converts biotin into the reactive intermediate biotin– AMP (adenosine monophosphate) to covalently label proteins proximal to TRIM21^30^ (Figure 4H). We treated cells with only biotin or biotin plus ACE, and then used streptavidin beads to capture biotinylated proteins. Label-free quantitative proteomics revealed that the ACE-treated group enriched NUP35, NUP54, NUP58, and GLE1 (Figure 4I). Notably, NUP35 and GLE1 were also the top depleted proteins following ACE treatment. These results, taken together, demonstrate that ACE induces the recruitment of TRIM21 to the vicinity of the NPC to trigger the degradation of nuclear pore proteins.

### TRIM21 is recruited to the nuclear pore via recognition of NUP98

As NPCs are the essential gateways connecting the nucleoplasm and cytoplasm, we wondered whether the degradation of nuclear pore proteins is responsible for ACE’s cytotoxicity. To answer this question, we aimed to identify mutations in the NPC that could render cells resistant to ACE. We exploited the recently described CRISPR-suppressor scanning method^31,32^ by designing a pool of tilling sgRNAs targeting the entire coding sequences of 27 nuclear pore proteins (omitting the nuclear ring-specific subunits), enabling the generation of large numbers of diverse nuclear pore protein variants in situ by the Cas9 nuclease (Figure 5A). A549-Cas9 cells transduced with the tilling sgRNA library were cultured in the absence or presence of ACE for three weeks. By next generation sequencing, we observed that sgRNAs targeting neighboring amino acids E721, K750, K752, and F777 in NUP98 were enriched in the ACE-treated group (Figure 5B).

**Figure 5:**
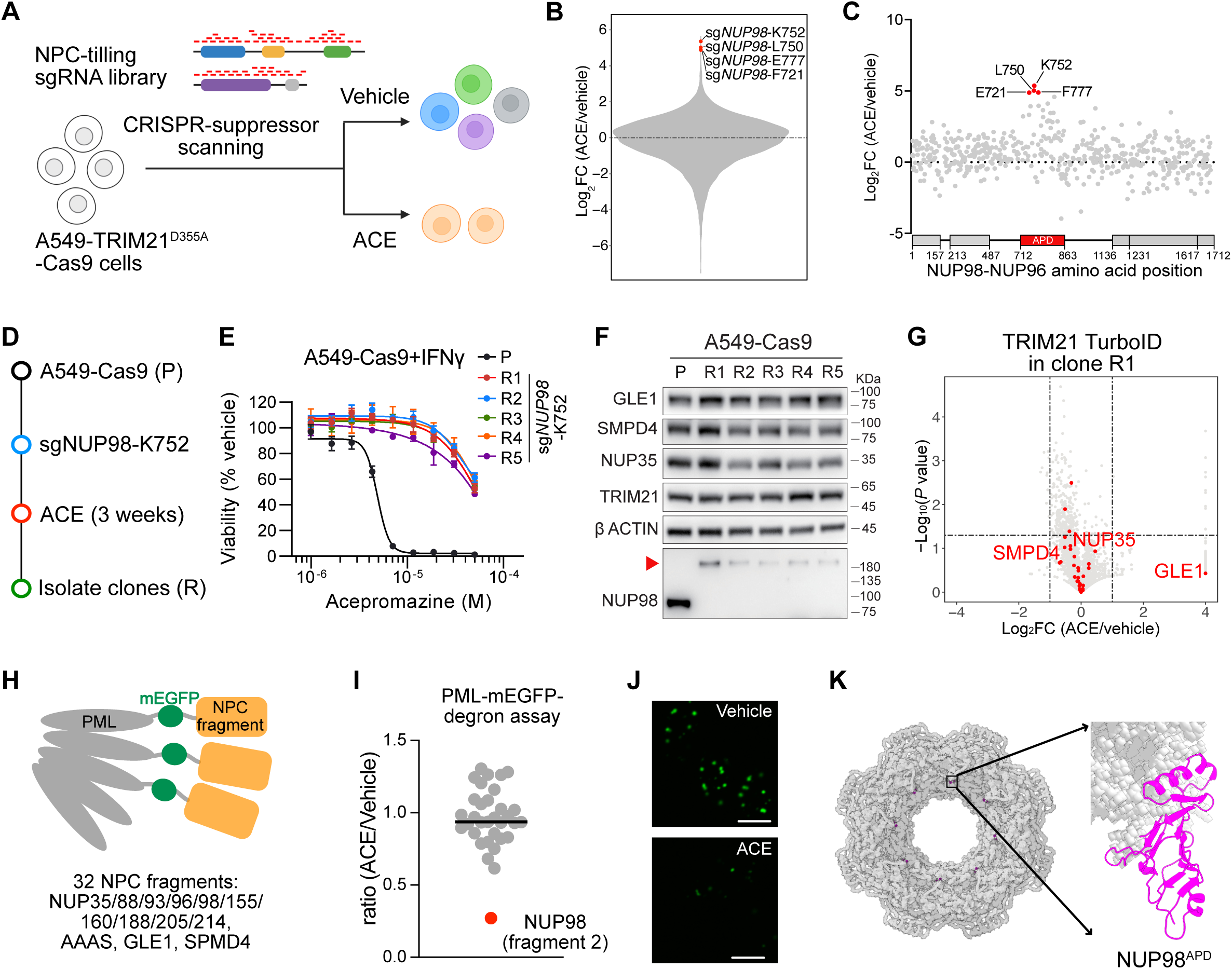
TRIM21 is recruited to the nuclear pore via recognition of NUP98. (A) Workflow of CRISPR-suppressor scanning (created with Biorender.com). (B) Violin plot showing log_2_ transformed fold change in the average abundance of each sgRNA comparing A549-TRIM21^D355A^ treated with ACE (dose escalating from 500 nM to 20 µM) versus vehicle for three weeks. Three independent samples were included in the analyses. Data used for the plots are provided in Table S6. (C) Scatter plot showing log_2_ transformed fold change in the average abundance of each NUP98-targeting sgRNA (y axis) in A549-TRIM21^D355A^ cells treated with ACE versus vehicle for three weeks. The sgRNAs are arrayed by the amino acid position in the NUP98-NUP96 coding sequence on the x axis corresponding to the position of the predicted cut site. Data points represent the mean values of three independent samples. (D) Workflow to isolate ACE-resistant clones from A549-Cas9 cells transduced with the sgRNA targeting NUP98 K752. (E) Concentration-response curves of ACE on the viability of *NUP98* mutant clones with IFNγ (10 ng/ml) pretreatment. Data indicate the mean ± s.e.m. of three independent samples. IC50 and 95% CI are shown in Table S1. (F) Immunoblots of indicated proteins in IFNγ-stimulated parental A549 cells and *NUP98* mutant clones (R1–R5) that were treated with ACE (10 μM) for 12 h. A representative result was shown from three independent experiments. Uncropped western blot images are provided as a Source Data file. (G) Volcano plot depicting log_2_ transformed average fold change of each protein (quantified by label-free proteomics) and −log_10_ transformed *P* value comparing TRIM21-TurboID enriched proteins from the *NUP98* mutant clone (R1) treated with ACE (20 µM) versus vehicle for 4 h. Three independent samples were included in the analyses. *P* values were calculated by unpaired Student’s t-test (two tailed). Data used for the plot are provided in Table S7. (H) Schematic of the PML-GFP-degron assay. (I) The effect of ACE on the levels of indicated PML-GFP-NPC fusion proteins in A549-TRIM21^D355A^ cells. Data are the mean ± s.e.m. of six independent samples. *P* values were calculated by unpaired Student’s t-test (two tailed). (J) Confocal microscopy images of PML-GFP-NUP98^domain^ ^2^ (green) in A549-TRIM21^D355A^ cells treat with vehicle or ACE (10 µM). Scale bar: 10 µm. (K) NUP98^APD^ in the cryo-electron tomography structure of the NPC (PDB: 7R5J).

NUP98 is a nuclear pore protein synthesized as a NUP98-NUP96 precursor, which undergoes autoproteolysis by its autoproteolytic domain (NUP98^APD^)^33^. The identified hotspot residues associated with ACE-resistance fell within NUP98^APD^ (Figure 5C). To further resolve the underlying mechanism of resistance, we transduced A549-Cas9 cells with the most enriched sgRNA (targeting K752) and selected five resistant clones following ACE treatment (Figures 5D-5E). Sequencing of NUP98 cDNA flanking the sgRNA cut site revealed the skipping of a short exon, resulting in the deletion of a segment (amino acids 733 to 770) in NUP98^APD^ (Figures S5A-S5B). Western blotting of these resistant clones revealed the disappearance of NUP98 and the appearance of a protein larger than 180 KDa, indicating the impairment of NUP98-NUP96 autoproteolytic processing (Figure 5F). By performing TRIM21-TurboID and quantitative proteomics assays, we found that ACE could no longer recruit TRIM21 to the vicinity of the NPC to degrade nuclear pore proteins in cells harboring mutant NUP98 (Figures 5G and S5C).

We explored how impaired NUP98-NUP96 processing could evade TRIM21 recognition to render ACE resistance. The complex organization of the NPC poses a challenge for detailed structure-function analysis. We thus designed a degron assay by taking advantage of the promyelocytic leukemia (PML) nuclear bodies, which are subnuclear structures formed via phase separation^34^. PML-GFP expression resulted in bright nuclear foci that could be easily visualized by high-content imaging. We then identified 13 proteins in the vicinity of NUP98 within the NPC and generated PML-GFP fusions of 32 fragments derived from these 13 proteins (Figure 5H). By high-content imaging, PML-GFP-NUP98^domain2^ stood out as most sensitive to ACE-mediated degradation (Figures 5H and 5I). By truncation analysis, we further narrowed down the minimal degron to amino acid positions 733-880 that resides in NUP98^APD^ (Figures 5I and S5D). Thus, CRISPR-suppressor screening and PML-GFP-degron assays converged to the same APD domain in NUP98, indicating that TRIM21 is recruited to the nuclear pore via recognition of NUP98^APD^.

### Crystal Structures of TRIM21^PRYSPRY^ in complex with acepromazine and its metabolites

To further explore the mechanism of action of ACE and its metabolites at the molecular level, we purified the PRYSPRY domain of TRIM21. Isothermal titration calorimetry (ITC) revealed modest binding affinities of ACE (dissociation constant/Kd: 5.66 µM), (*R*)-ACE-OH (Kd: 9.11 µM), and (*S*)-ACE-OH (Kd: 17.9 µM) to TRIM21^PRYSPRY^(D355A) (Figure 6A). No binding signal was detected with WT TRIM21^PRYSPRY^ up to 50 µM of titrated compounds. We then used microscale thermophoresis (MST) to complement ITC findings and observed weak binding of ACE (Kd: 44.6 µM) and (*S*)-ACE-OH (Kd: 237.3 µM) to WT TRIM21^PRYSPRY^ (Figure S6A). No binding of (*R*)-ACE-OH to WT TRIM21^PRYSPRY^ could be detected by MST. These biophysical measurements revealed weak binding of ACE and its metabolites to TRIM21^PRYSPRY^, which alone could not explain why (*S*)-ACE-OH was active whereas ACE and (*R*)-ACE-OH were inactive.

**Figure 6:**
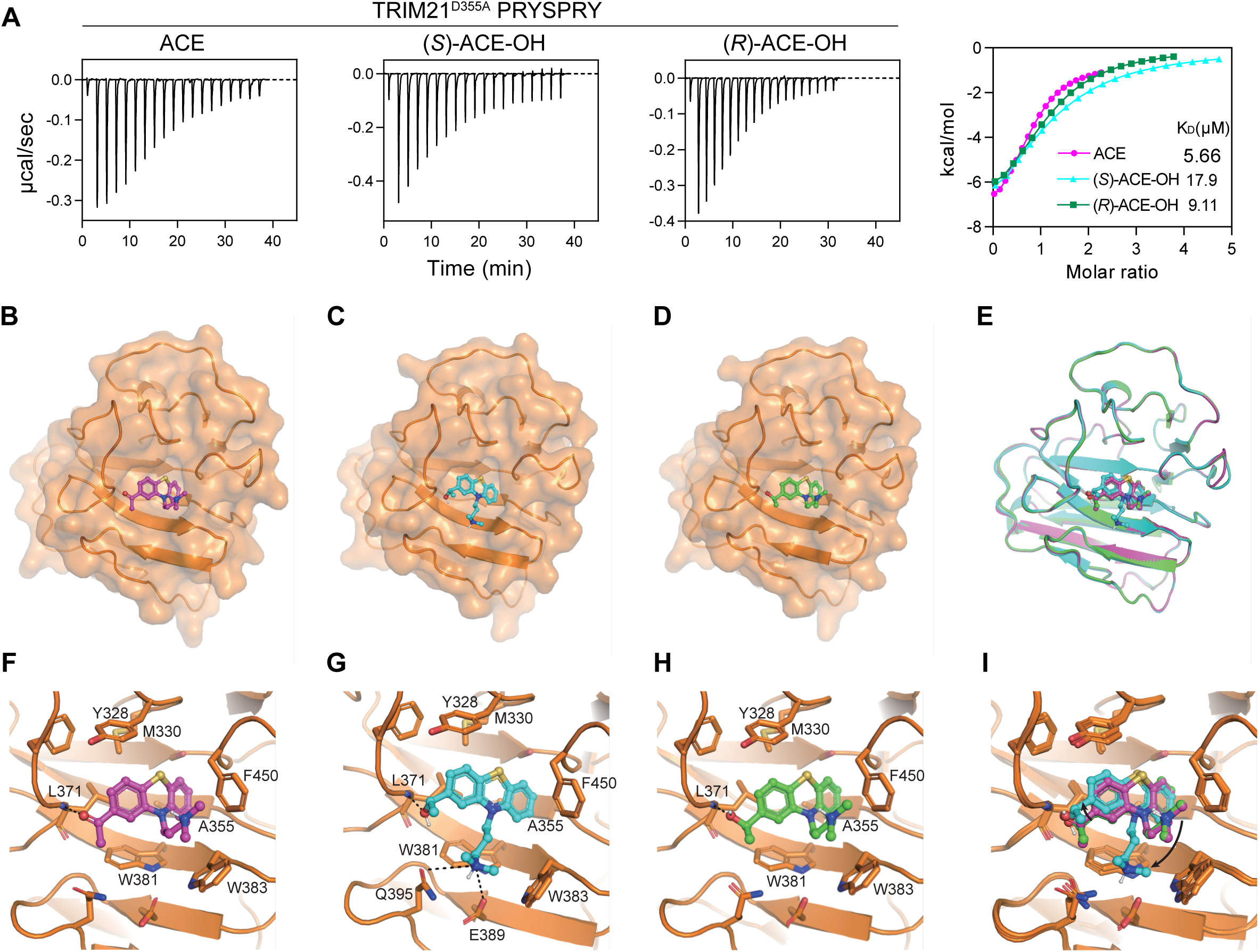
Crystal Structures of the PRYSPRY domain of TRIM21^D355A^ in complex with acepromazine and its metabolites. (A) ITC titration and curve fitting of ACE and its metabolites with the PRYSPRY domain of TRIM21^D355A^ (B-D) The overall structure of the PRYSPRY domain of TRIM21^D355A^ in complex with ACE (B), (*S*)-ACE-OH (C), and (*R*)-ACE-OH (D). Crystallographic data are provided in Table S9. (E) Superimposed binding positions for ACE (magenta), (*S*)-ACE-OH (cyan), and (*R*)-ACE-OH (green) within the PRYSPRY domain of TRIM21^D355A^. (F) - (H) Close-up views of the binding poses for ACE (F), (*S*)-ACE-OH (G), and (*R*)-ACE-OH (H) within the pocket of the PRYSPRY domain of TRIM21^D355A^. (I) Superimposed ligand conformations of ACE (magenta), (*S*)-ACE-OH (cyan) and (*R*)-ACE-OH (green).

We next determined the co-crystal structures of TRIM21^PRYSPRY^(D355A) in complex with ACE, (*S*)-ACE-OH and (*R*)-ACE-OH with resolutions of 1.6 Å, 1.89 Å and 1.74 Å, respectively (Figures 6B-6E and S6B). These structures revealed that the tricyclic phenothiazine rings of these ligands occupy a shallow hydrophobic pocket formed by Y328, M330, A335, L371, W383, and F450. The oxygen atom in the acetyl or hydroxyl group of the ligands acts as a hydrogen bond acceptor, interacting with the backbone of L371 (Figures 6F-6H). Notably, the aliphatic chain of the ligands exhibits flexibility, resulting in different orientations of the amine group. In the case of ACE and (*R*)-ACE-OH, the amine group extends outside the pocket, whereas in the case of (*S*)-ACE-OH, the amine group forms polar interactions with the side chains of E389 and Q395 (Figure 6I). As (*S*)-ACE-OH is the active metabolite, these structural observations suggest that the side chain conformation of these ligands could be a critical determinant of molecular glue activity.

### TRIM21-based PROTACs selectively degrade multimeric proteins in biomolecular condensates

The co-crystal structures of TRIM21^PRYSPRY^(D355A) in complex with ACE revealed the aliphatic chain as a potential exit vector for PROTAC design. We thus generated a TRIM21-based PROTAC (TrimTAC1) by linking ACE and JQ1 (a high affinity binder of BRD4) with a three-carbon aliphatic chain (Figure 7A). As a positive control, we used dBET1, which is a CRBN-based PROTAC targeting BRD4^35^. To provide cellular models for evaluating PROTAC activity, we stably expressed mEGFP fused to the second bromodomain of BRD4 (BRD4^BD2^) in A549 cells expressing TRIM21(D355A) or CRBN. While dBET1 potently (half-maximal degradation concentration/DC_50_: 31 nM) degraded mEGFP-BRD4^BD2^, TrimTAC1 could not degrade mEGFP-BRD4^BD2^ at concentrations up to 100 µM (Figures 7B-7C).

**Figure 7:**
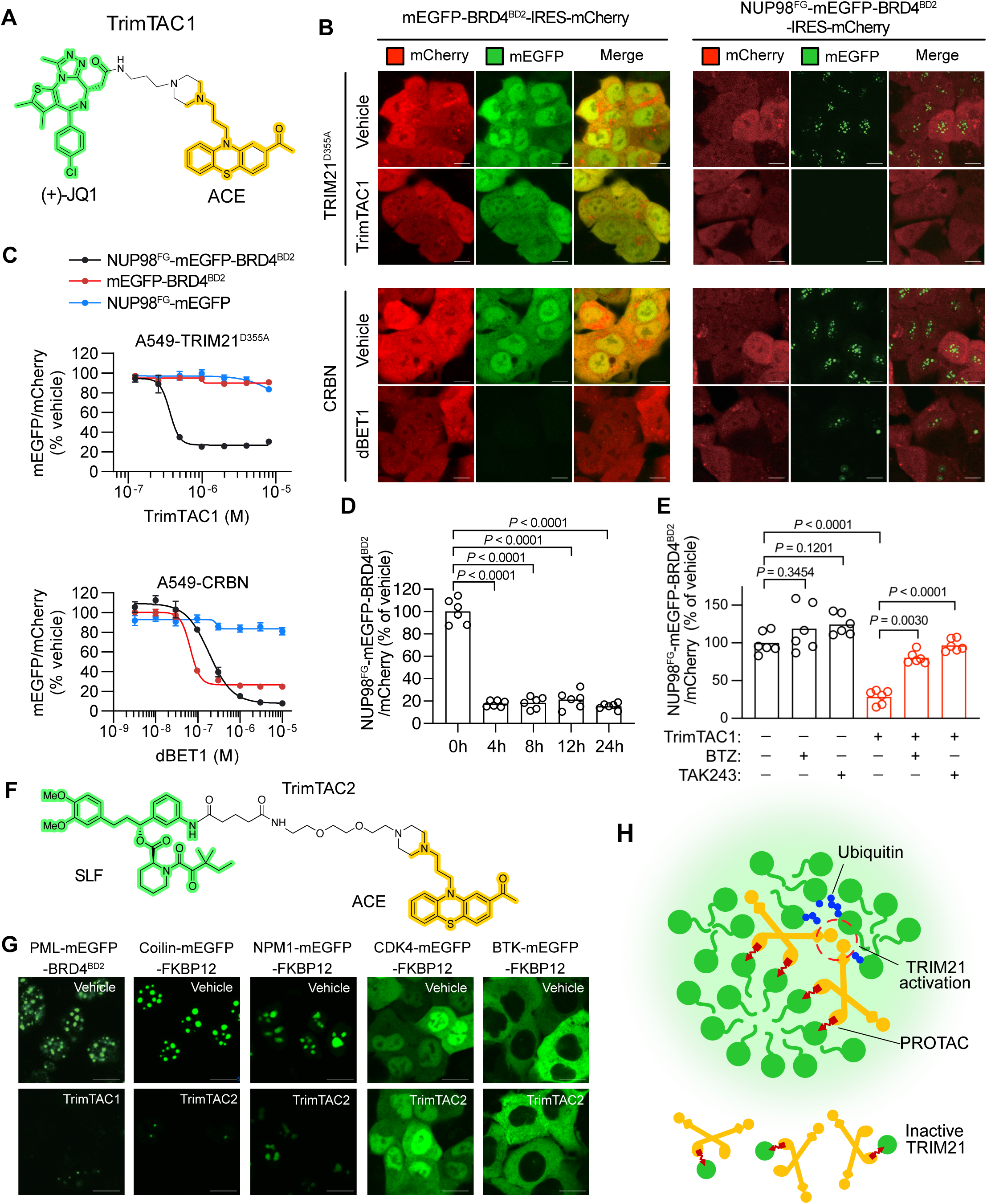
TRIM21-directed PROTACs selectively degrade multimeric proteins in biomolecular condensates. (A) Chemical structure of TrimTAC1. (B) Representative images of A549-TRIM21^D355A^ or A549-CRBN cells expressing indicated reporter proteins treated with or without TrimTAC1 (4 µM) or dBET1 (100 nM) for 4 h. Scale bar: 10 µM. (C) Concentration-response curves of TrimTAC1 or dBET1 on the normalized intensities of indicated reporter proteins in A549-TRIM21^D355A^ or A549-CRBN cells. Data indicate the mean ± s.e.m. of three independent samples. IC50 and 95% CI are shown in Table S1. (D) Intensities of NUP98^FG^-mEGFP-BRD4^BD2^ in A549-TRIM21^D355A^ cells treated with TrimTAC1 (2 µM) for indicated time. Each point represents the mean of three independent samples. One-way ANOVA followed by Dunnett’s multiple comparison test was used to determine statistical significance. (E) Normalized intensities of NUP98^FG^-mEGFP-BRD4^BD2^ in A549-TRIM21^D355A^ cells with indicated treatment of TrimTAC1 (4 µM), BTZ (400 nM), and TAK243 (1.5 µM). Data indicate the mean ± s.e.m. of six independent samples. One way ANOVA followed by Tukey’s multiple comparison test was used to determine statistical significance. (F) Chemical structure of TrimTAC2. (G) Representative images of indicated FKBP12-mEGFP reporter proteins treated with vehicle, TrimTAC1 (10 μM), or TrimTAC2 (10 μM) for 4 h. Scale bar: 10 µm. (H) Schematic illustration of TrimTAC selectivity towards multimeric proteins in biomolecular condensates.

TRIM21 has been exploited in the Trim-Away technology, which uses antibodies to direct TRIM21 to degrade intracellular proteins^36^. A recent mechanistic study revealed that substrate-induced clustering triggers intermolecular dimerization of the RING domains of TRIM21 to switch on its enzymatic activity^37^. Interestingly, NUP98^APD^, the degron recognized by TRIM21, is also embedded in the multimeric nuclear pore complex (Figure 5K). We therefore hypothesized that TRIM21-based PROTACs could drive selective degradation of target proteins present in multimeric assemblies that permits the dimerization of TRIM21 RING domains.

To test this hypothesis, we fused the N-terminal FG (phenylalanine-glycine) repeat domain of NUP98 to the mEGFP-BRD4^BD2^ reporter. The resulting fusion protein (NUP98^FG^-mEGFP-BRD4^BD2^) formed nuclear condensates in A549 cells, which were efficiently and rapidly degraded by both TrimTAC1 and dBET1 when TRIM21(D355A) or CRBN were respectively expressed (Figures 7B-7D). Pretreating cells with inhibitors targeting the proteasome (bortezomib) or the E1 ubiquitin activating enzyme (TAK243) prevented TrimTAC1-induced degradation of NUP98^FG^-mEGFP-BRD4^BD2^ nuclear condensates (Figures 7E and S7A). To avoid over-reliance on JQ1, we synthesized TrimTAC2 using the SLF warhead (Synthetic Ligand of FKBP12)^38^ (Figure 7F). TrimTAC2 efficiently degraded the NUP98^FG^-mEGFP-FKBP12 condensates without affecting the soluble mEGFP-FKBP12 (Figures S7B-S7E).

To further test the generalizability of TrimTACs to degrade proteins present in biomolecular condensates, we fused mEGFP-FKBP12 to three additional condensate-forming proteins: PML (PML-NB), Coilin (Cajal bodies), and NPM1 (nucleolus)^39–41^. As examples of non-condensate proteins, we fused mEGFP-FKBP12 to the kinases CDK4 and BTK. In A549 cells expressing TRIM21(D355A), TrimTAC2 degraded condensate-forming fusion proteins (PML, Coilin, and NPM1), but did not degrade fusion proteins that do not form condensates (CDK4 and BTK) (Figure 7G). Taken together, these results indicate that TRIM21-based PROTACs possess the ability to degrade proteins in biomolecular condensates and spare proteins in the dilute phase.

## Discussion

As a mechanism of intracellular immunity, TRIM21 uses antibodies as bridging molecules to direct invading pathogens for proteasomal degradation^26^. This capability of TRIM21 was recently exploited as the Trim-Away technology to enable acute and rapid degradation of endogenous cellular proteins^36^. Despite its enormous utility, Trim-Away requires electroporation or microinjection to deliver antibodies to cells, which is currently not feasible in most therapeutic settings. TRIM21-based degraders reported here provide chemical starting points to extend TRIM-Away to therapeutic applications.

TRIM21 forms inactive dimers in which a central antiparallel coiled coil positions the two RING domains at the two ends of the dimer. Upon binding to multimeric substrates, RING domains from neighboring TRIM21 dimerizes, leading to the activation of its enzymatic activity^37^. In line with substrate-induced activation of TRIM21, we present two classes of TRIM21-based degraders with a high degree of selectivity towards multimeric neo-substrates. The first class, represented by the serendipitously discovered (*S*)-ACE-OH, functions as a monovalent degrader to guide TRIM21 to recognize NUP98, resulting in the degradation of multiple proteins in the nuclear pore complex. The second class, represented by the rationally designed bivalent TrimTACs, degrades multimeric proteins localized to various biomolecular condensates without affecting monomeric proteins in the dilute phase (Figure 7H). These results highlight the potential of TRIM21-based degraders to differentially eliminate aberrant proteins in multimeric assemblies leading to neurodegeneration, cancer, and autoimmunity^42–44^. Such multimer-selective degraders would be particularly valuable in situations where the degradation of monomeric proteins could cause undesirable consequences.

Our structural biology efforts revealed that TRIM21-based degraders exploit a ligandable pocket in the C-terminal PRYSPRY domain of TRIM21, which features six extended loops that are analogous to the CDR (complementarity-determining region) loops in antibodies^28^. The ligandable pocket and the flexibility of CDR loops in TRIM21^PRYSPRY^ could be harnessed in the future to create neomorphic interfaces for molecular glues to target other disease-causing proteins. PRY and SPRY domains are the most common C-terminal domains in TRIM family members^25,45^. The SPRY domain is found alone in 39 human TRIM family members, and is fused to the PRY domain (to form the PRYSPRY domain) in 24 family members. The extent to which these domains can be engaged by bifunctional, or even molecular glue degraders to promote targeted protein degradation should represent a fertile area for future investigation.

## Methods

### Cell culture

A549, HCT-116, HeLa, DLD-1, and HEK293T were gifts from Dr. Deepak Nijhawan’s lab at University of Texas Southwestern Medical Center. NK-92MI was a gift from Dr. Feng Shao’s lab at National Institute of Biological Sciences, Beijing. Mino was obtained from ATCC. SiHa, HuH-7, ME-180, and HaCaT were obtained from Cell Resource Center, Peking Union Medical College (Beijing, China). Activated human T lymphocytes from healthy donors were gifts from Yanping Ding at Immunochina Pharmaceuticals Co., Ltd., Beijing, China. All cell lines were confirmed to be mycoplasma free by PCR. Regular cell culture methods were used to culture cells in tissue-culture incubators with 5% CO2 at 37°C. A549 and DLD-1 were grown in RPMI-1640 medium with 10% fetal bovine serum (FBS) and 2 mM L-glutamine. HCT-116, HEK293T, HeLa, SiHa, HuH-7, ME-180, and HaCaT cells were grown in DMEM medium with 10% FBS and 2 mM L-glutamine. NK-92MI cells were grown in alpha Minimum Essential Medium (MEM) supplemented with 0.2 mM inositol, 0.1 mM 2-mercaptoethanol, 0.02 mM folic acid, 10mM HEPEs, non-essential amino acid (NEAA), and 20% FBS.

### Chemicals

Acepromazine (CAS: 3598-37-6), chlorpromazine (CAS: 69-09-0), bortezomib (CAS: 179324-69-7), MLN4924 (CAS: 905579-51-3), TAK243 (CAS: 1450833-55-2), biotin (CAS: 58-85-5) were purchased from TargetMol (Topscience, Shanghai, China). Chemicals were prepared as 10 mM stocks in dimethyl sulfoxide (DMSO, CAS: 67-68-5, Solarbio Life Science, Beijing, China) and further diluted in DMSO to the desirable concentrations.

### High-throughput small-molecule screening

A compound library with 81,845 small molecules (Life Chemicals, Niagara-on-the-Lake, Canada) was used in high-throughput screening. WT A549 cells (labeled with H2B-GFP) and *IFNGR1*-deficient A549 cells (labeled with H2B-mCherry) were mixed at 1:1 ratio. Three thousand mixed cells in 50 μL of medium were plated per well in 384-well flat clear bottom white polystyrene TC-treated microplates (Corning, Corning, USA) and allowed to attach to plates overnight. Compounds from the screening library were added to the cells with a Biomek FXP automated workstation at a final concentration of 10 μM. Four days later, images were collected with a PerkinElmer Opera LX high-content screening system. The ratios of GFP^+^ nuclei to mCherry^+^ nuclei were measured by the Columbus software.

### NK killing assay

Ten thousand A549 cells were seeded per well in 6-well plates and allowed to attach to the plates overnight. NK-92MI cells were added at various effector/target (E/T) ratios together with DMSO or 10μM acepromazine. Two days later, cells were stained with crystal violet (Beyotime, Shanghai, China, C0121) for 20 min at room temperature, and then de-stained with water. Images were taken with a VILBER FX7 imager. To perform the quantitative NK killing assay, WT A549 cells (labeled with H2B-GFP) and *IFNGR1*-deficient A549 cells (labeled with H2B-mCherry) were mixed at 1:1 ratio and co-incubated with different amounts of NK-92MI cells in the presence or absence of 10μM acepromazine. Two days later, the ratios of GFP^+^ cells to mCherry^+^ cells were measured by a LSR Fortessa flow cytometer (BD Biosciences, Franklin Lakes, NJ, USA).

### Cell viability assay

Three thousand A549 cells in 100 μL of medium were plated per well in 96-well flat clear bottom white polystyrene TC-treated microplates (Corning, Corning, USA). Cells were dosed with a serial dilution of compounds with a D300e digital dispenser (Tecan, Männedorf, Switzerland). Cell survival was measured three days later using CellTiter-Glo luminescent cell viability assay kit (Promega, Madison, WI, USA) according to the manufacturer’s instructions. Luminescence was recorded by EnVison multimode plate reader (PerkinElmer, Waltham, MA, USA). IC50 was determined with GraphPad Prism using baseline correction (by normalizing to the DMSO control), the asymmetric (four parameters) equation, and least squares fit.

### Metabolite extraction and LC-MS analysis

A549 or HCT-116 cells cultured in 6-well plates were treated with 10 μM ACE for 24 h. The conditioned medium was collected and ethyl acetate were added at a 2:1 ratio. The mixture was vortexed for 30s and centrifuged for 5 min at 15000 rpm. The metabolites were extracted into the upper layer of ethyl acetate. LC-MS analysis was performed on a Waters system (Column: BEH C18, 1.7 µm, 2.1*50 mm) with a Photodiode Array (PDA) detector and a Single Quadrupole (SQ) detector.

### Genome-wide CRISPR screening

A549-Cas9 cells were infected with the Brunello CRISPR sgRNA library^22^ at a multiplicity of infection (MOI) of ∼0.3 and a coverage of ∼100 cells per sgRNA. Twenty-four hours later, the infected cells were re-seeded and selected with 2 µg/ml puromycin for 3 days. Cells surviving puromycin selection were treated with 10 ng/ml IFNγ together with DMSO or ACE (gradually increasing from 4 µM to 10 µM) for four weeks to select for resistant cells. Extraction of genomic DNA and amplification of sgRNA for next generation sequencing were performed as described^46^. CRISPR screening data were analyzed by MAGeCK (v0.5.9.2)^23^.

### Western blotting

Cells were washed with DPBS to remove residual medium and then lysed in SDS lysis buffer (20 mM HEPES, 2 mM MgCl_2_, 10 mM NaCl, 1% SDS, pH 8.0) containing 0.5 units/µL benzonase, EDTA-free protease inhibitor cocktail (Roche, Basel, Switzerland). The protein lysates were centrifuged (12000 g, 10 min, 4 °C), and the concentration of the supernatants was determined using the BCA Protein Assay. Proteins were separated on a 4%-20% gradient SDS–PAGE gel and transferred to nitrocellulose membranes with a pore size of 0.5 microns. The membranes were blocked in 5% nonfat milk PBST solution (0.1% v/v Tween-20) for 30 min and then were sequentially incubated with the primary antibody overnight at 4 °C and the secondary antibody at room temperature for 1 h. The primary antibodies used are as follows: rabbit polyclonal anti-Stat1 (1:5,000, 9172, Cell Signaling Technology, Danvers, MI, USA), rabbit monoclonal anti-Phospho-Stat1 (1:5,000, 7649, Cell Signaling Technology, Danvers, MI, USA), mouse monoclonal anti-Flag-HRP (1:10000, A8592, Sigma-Aldrich, St. Louis, MO, USA), mouse monoclonal anti-β-actin-HRP (1:10,000, HX18271, Huaxingbio, Beijing, China), rabbit monoclonal anti-TRIM21 (1:5000, AB207728, Abcam, Cambridge, UK), rabbit polyclonal anti-GLE1 (1:5000, A13207, ABclonal, Woburn, MA, USA), anti-SMPD4 (1:5000, A15473, ABclonal, Woburn, MA, USA), anti-NUP35 (1:5000, A12762, ABclonal, Woburn, MA, USA), and anti-NUP98 (1:5000, A0530, ABclonal, Woburn, MA, USA). The secondary antibody used is goat polyclonal anti-rabbit IgG (1:10,000, 7074, Cell Signaling Technology, Danvers, MI, USA). M5 HiPer ECL Western HRP Substrate (Mei5bio, Beijing, China) was used for the detection of HRP enzymatic activity. Western blot images were taken with a VILBER FUSION FX7 imager.

### Quantitative mass spectrometry

Cell pellets were lysed with lysis buffer (8 M urea, 50 mM Tris-HCl, 1% Triton X-100, pH=7.4) containing protease and phosphatase inhibitors (539134 and 524625, Merck, Rahway, NJ, USA). Cell lysates were reduced with 10 mM dithiothreitol (DTT), alkylated with 40 mM iodoacetamide, then quenched with 5 mM DTT. Alkylated samples were cleaned up using the SP3 method followed by trypsin digestion as previously described^47^. All DIA-MS experiments were performed on an Orbitrap Exploris 480 equipped with an UltiMate 3000_UPLC system (Thermo Scientific, Waltham, MA, USA). The project-specific DIA spectral library was first generated by DIA-MS2pep as previously described^48^. Protein sequences for database search was reviewed Human proteome (uniport _ UP000005640, 82685 entries). The run-specific FDR of identifications at both peptide and protein level were estimated by Percolator^49^. The spectral library was further generated and submitted to DIA-NN software^50^ for protein quantification. Precursor and protein FDR was set to 1%.

### TurboID

A549-TRIM21-TurboID cells were treated with 10 ng/ml IFNγ overnight, 100 nM bortezomib for 8 hours, vehicle (DMSO) or 20 μM ACE for 6 hours, and 50 μM biotin for 2 hours. Cells were scraped off plates, pelleted by centrifugation, and rinsed with PBS. Cell pellets were lysed in the RIPA lysis buffer (50 mM Tris pH 8, 150 mM NaCl, 1%SDS, 0.5% sodium deoxycholate, 1% Triton X-100, 1× protease inhibitor cocktail (Sigma-Aldrich, St. Louis, MO, USA), and 1 mM PMSF) and incubated with streptavidin magnetic beads (88817, Thermo Scentific, Waltham, MA, USA) at 4℃ overnight. Beads were sequentially washed with 1 M KCl, 0.1 M Na_2_CO_3_, RIPA buffer, and TBS (50 mM Tris-HCl pH 7.5, 150 mM NaCl). Bound proteins were eluted with elution buffer 1 (50 mM Tris-HCl pH 7.5, 2 M urea, 5 ng/µL sequencing grade modified trypsin (Promega, Madison, WI, USA), 1 mM DTT) and elution buffer 2 (50 mM Tris-HCl pH 7.5, 2 M urea, 5 mM iodoacetamide). The eluted proteins were digested overnight at 400 rpm, 32°C followed by acidification with trifluoroacetic acid and drying by vacuum centrifugation. LC-MS/MS experiments were performed on an Orbitrap Exploris 480 equipped with an UltiMate 3000_UPLC system (Thermo Scientific, Waltham, MA, USA) as described above.

### CRISPR-suppressor screening

The NPC tiling sgRNA library included every sgRNA with an NGG PAM and a cleavage site within the coding sequences of 27 nuclear pore proteins. The oligo pool containing the sgRNA sequences was synthesized by Genewiz (Suzhou, Jiangsu, China) and cloned into lenti-guide-puro (#52963; Addgene, Watertown, MA, USA) as previously described^51^. Lentiviral particles carrying the resultant NPC tiling library were generated and titered as previously described^51^. A549-TRIM21(D355A)-Cas9 cells were infected with the NPC tiling sgRNA library at a multiplicity of infection (MOI) of ∼0.3 and a coverage of ∼1000 cells per sgRNA. After selection with 2 µg/ml puromycin for three days, cells were treated with DMSO or ACE (gradually increasing from 0.5 µM to 20 µM) for three weeks. Extraction of genomic DNA and amplification of sgRNA for next generation sequencing were performed as described^46^.

### Isolation of acepromazine-resistant A549 clones

A549-Cas9 cells were infected with lentivirus harboring sg*Nup98*-K752. Stable cell lines were obtained by selection of transduced cells with 2 µg/ml of puromycin. Afterwards, cells were treated with 10 μM acepromazine continuously for three weeks. Acepromazine-resistant clones were isolated, expanded, and tested for their sensitivity to acepromazine.

### Protein expression and purification

Human TRIM21 PRYSPRY (residues 287-465) was cloned into the pET28a vector with an N-terminal 6xhis tag. The D355A mutation was introduced by overlap extension PCR. Bacterial protein expression was carried out in E. coli BL21-CodonPlus(DE3)-RIL cells. Cells were grown in 2xTY medium (supplemented with 0.5% glucose, 2 mM MgSO_4_ and Kanamycin) at 37 °C and induced at OD600 around 0.6–1 with 1 mM isopropyl-β-D-thiogalactopyranoside (IPTG). Cells were lysed by sonication in buffer A (50 mM Tris-HCl pH 8.0, 1M NaCl, 2 mM TECP) supplemented with 10 mM imidazole and 1x cOmplete, Mini, EDTA-free protease inhibitor cocktail (Roche, Bazel, Switzerland). The lysate was clarified by centrifugation at 15,000 g for 1 hour at 4 °C. Recombinant proteins were purified by Ni NTA Beads (SA004010, Smart Lifesciences, Changzhou, China) and washed with buffer B (50 mM Tris-HCl pH 8.0, 300 mM NaCl, 1 mM TECP, 30 mM imidazole). Proteins were eluted with buffer C (50 mM Tris-HCl pH 8.0, 300 mM NaCl, 1 mM TECP, 30 mM imidazole). The flow-through containing TRIM21 PRYSPRY was concentration by ultrafiltration, loaded onto Superdex 75 Increase 10/300 GL column (Cytiva, Marlborough, MA, USA), and then fractionated in 50 mM Tris-HCl pH 8.0, 150 mM NaCl, 0.5 mM TECP. Purified proteins were concentrated to 20 mg/mL by ultrafiltration.

### Isothermal titration calorimetry

ITC was performed using a Microcal PEAQ-ITC (Malvern Panalytical, Malvern, Worcestershire, United Kingdom) with protein solution in the cell and compound solution in the syringe. Experiments were carried out at 25 °C. The titrations consisted of 19 injections, an initial injection of 0.4 μL and then 18 injections of 2 μL. The stirring speed in the reaction cell was 1000 rpm. Data were analyzed using non-linear least squares regression using Microcal PEAQ analysis software. Drifts in the baseline were corrected during data analysis.

### Crystallization and X-ray data collection

Co-crystals were obtained by mixing ACE and its metabolites with purified TRIM21 PRYSPRY(D355A) at a molar ratio of 2:1. All crystals were grown by a vapor diffusion method in hanging-drops at 20 °C. Hampton Crystal Index were used for initial crystallization screens. The best co-crystals were grown at 20 °C above a well solution containing 0.1 M Tris (pH 7.0), 3.5 M sodium formate. Macroseeding was used to improve crystal quality. X-ray diffraction data were collected at Shanghai Synchrotron Radiation Facility BL02U1 beamline and BL17UM beamline^52,53^. Data were processed using the XDS, POINTLESS and AIMLESS programs^54–56^.

### Structure determination and Refinement

The crystal structure of TRIM21 in complex with ligands were solved by molecular replacement with the program Phaser^57^, using a previous reported TRIM21-IgG-Fc complex structure (PDB ID: 2IWG)^28^ as the search model. Further manual model building was facilitated by using Coot^58^, combined with the structure refinement using phenix.refine^59^.

### MST measurements between TRIM21 PRYSPRY and compounds

Purified TRIM21 PRYSPRY domain was labeled using Protein Labeling Kit RED-NHS 2nd Generation (MO-L011, NanoTemper Technologies GmbH, Munich, Germany). Labeled protein was diluted to 50 nM in PBST (137 mM NaCl, 2.5 mM KCl, 10 mM Na2HPO_4_, 2 mM KH2PO_4_, pH 7.4, 0.05% Tween-20) during measurement. ACE, (*R*)-ACE-OH, and (*S*)-ACE-OH were prepared by serial dilution in PBST. Protein and compound mixtures were loaded into standard treated capillaries (MO-K022, NanoTemper Technologies GmbH, Munich, Germany). Measurements were performed on a Monolith NT.115 with blue/red filters (NanoTemper Technologies GmbH, Munich, Germany) at 25 °C using 80% MST power. All experiments were repeated three times for each measurement. Data analyses were performed using the NanoTemper® Affinity Analysis software.

### Confocal imaging

Cells expressing fluorescent proteins were cultured on glass bottom plates (801002, NEST, Wuxi, Jiangsu, China) and stained with Hoechst (1 µg/ml, abs813337, Absin Biosciences, Shanghai, China) for 5 minutes before capturing. All confocal images were captured using a Nikon A1 sim confocal microscope and were then processed using IMARIS software.

### Transmission electron microscopy imaging

Cells grown on sapphire discs were treated with 10 ng/ml IFNγ and 10 μM ACE and cryoimmobilized by high pressure freezing (HPF COMPACT 01, Engineering office M. Wohlwend; Sennwald, Switzerland) with 0.25 mm-deep HPF carriers as caps and 1-hexadecene as a cryoprotectant. The frozen samples were maintained under liquid nitrogen before freeze-substitution. For optimal ultrastructural preservation, samples were transferred to the freeze-substitution medium (acetone containing 1% osmium tetroxide, 0.1% uranyl acetate, and 5% H2O) under liquid nitrogen and placed in an automatic freeze substitution system (Leica AFS2, Leica Microsystems, precooled to −90 ℃) for 24 h. The samples were subsequently warmed to −60 ℃ over 6 h, kept at −60 ℃ for 8 h, and warmed to −30 ℃ over 6 h, kept at −30 ℃ for 8 h. Then the samples were warmed to 4 ℃ over 3 h, and kept at 4 ℃ for 15 min before washing with anhydrous acetone. Samples were then infiltrated and embedded in SPI-Pon 812 resin. Ultrathin sections (90 nm-thick) were cut with an ultramicrotome (Leica EM UC7, Leica Microsystems), and stained with 3% uranyl acetate in 70% methanol/H2O for 7 min, followed by Sato’s lead for 2 min. Images were obtained on a TECNAI spirit G2 (FEI; Eindhoven, Netherlands) transmission electron microscope at 120 kV.

### High-content imaging

For high-content imaging, cells expressing fluorescent proteins (mEGFP and mCherry) were plated in 384-well flat clear bottom white polystyrene TC-treated microplates (Corning, Corning, USA) and allowed to attach to plates overnight. After compound treatment, images were collected with a PerkinElmer Opera LX high-content screening system. The total intensity of EGFP and mCherry were measured by the Columbus software.

### Reporter protein degradation assay

For the PML degron assay, 32 fragments from 13 NPC proteins were fused at the C-terminus with PML-mEGFP, and stably expressed in A549-TRIM21(D355A) cells via lentiviral transduction. For the TrimTAC assays, NUP98^FG^, PML, Coilin, NPM1, CDK4, and BTK were fused at the N-terminus with mEGFP-BRD4^BD2^ or mEGFP-FKBP12, and stably expressed in A549-TRIM21(D355A) or A549-CRBN cells via lentiviral transduction. Compound-induced degradation of reporter proteins was analyzed by high-content imaging.

### Quantification and statistical analysis

Statistical analyses were performed with GraphPad Prism 8.0. Student’s t test was used to evaluate the statistically significant difference between the two sample groups. When comparing more than two independent groups, ANOVA was used to evaluate statistical significance. Multiple comparison tests were performed when ANOVA was significant. All tests were two-tailed, and *P* < 0.05 was considered statistically significant.

## Data availability

The co-crystal structures of TRIM21(D355A)^PRYSRY^ has been deposited in PDB (8Y58, 8Y59, 8Y5B). Data are available within the Article, Supplementary Information or Source Data file. Source data are provided with this paper.

## Code availability

The following open-source code and software were used in this study: MAGeCK (v0.5.9.4). The following R libraries were used: R (v4.1.2), RStudio (v2021.9.2.382), readxl (v1.3.1), writexl (v1.4.2), tidyverse (v1.3.1), MAGeCKFlute (v1.14.0), latex2exp (0.9.6). Other software: Prism (v8.0.2), Affinity Designer (v2.2.0), ChemBioDraw Ultra (v14.0), UCSF-Chimera (v1.16).

## Acknowledgements

We thank the staff at BL02U1 and BL17UM beamlines at SSRF for help in X-ray diffraction data collection and analysis, Cheng Chen for synthesizing TrimTACs, Wei Wang for help with crystal screening, Pilong Li for sharing plasmids and helpful discussions, NIBS imaging, TEM, and metabolomics core facilities for technical support. This work was supported by Beijing Municipal Commission of Science and Technology (Z201100005320010 and Z221100003422015 to T.H.; Z221100007022004 to N.H.), funding from National Institute of Biological Sciences, Beijing and Tsinghua Institute of Multidisciplinary Biomedical Research (to T.H. and N.H.).

## Author contributions

T.H., P.L., Y.C., and L.X. conceived ideas, designed experiments, and wrote the paper. P.L., Y.C., and L.X. conducted most of the experiments, analyzed the data, and prepared the figures. X.R. solved all co-crystal structures. Q.W., C.C., and C.L. identified the chirality of the active ACE-OH enantiomer. J.L. and X.Q. helped with high-throughput screening. J.H. and W.J. performed quantitative mass spectrometry, S.S. helped with the PML degron assay. T.H. and N.H. obtained funding and supervised the study.

## Competing interest declaration

A provisional patent application has been filed. The authors declare no competing interest.

## Additional information

Correspondence and requests for materials should be addressed to Niu Huang or Ting Han.

## Supplementary Figure legends

**Figure S1:**
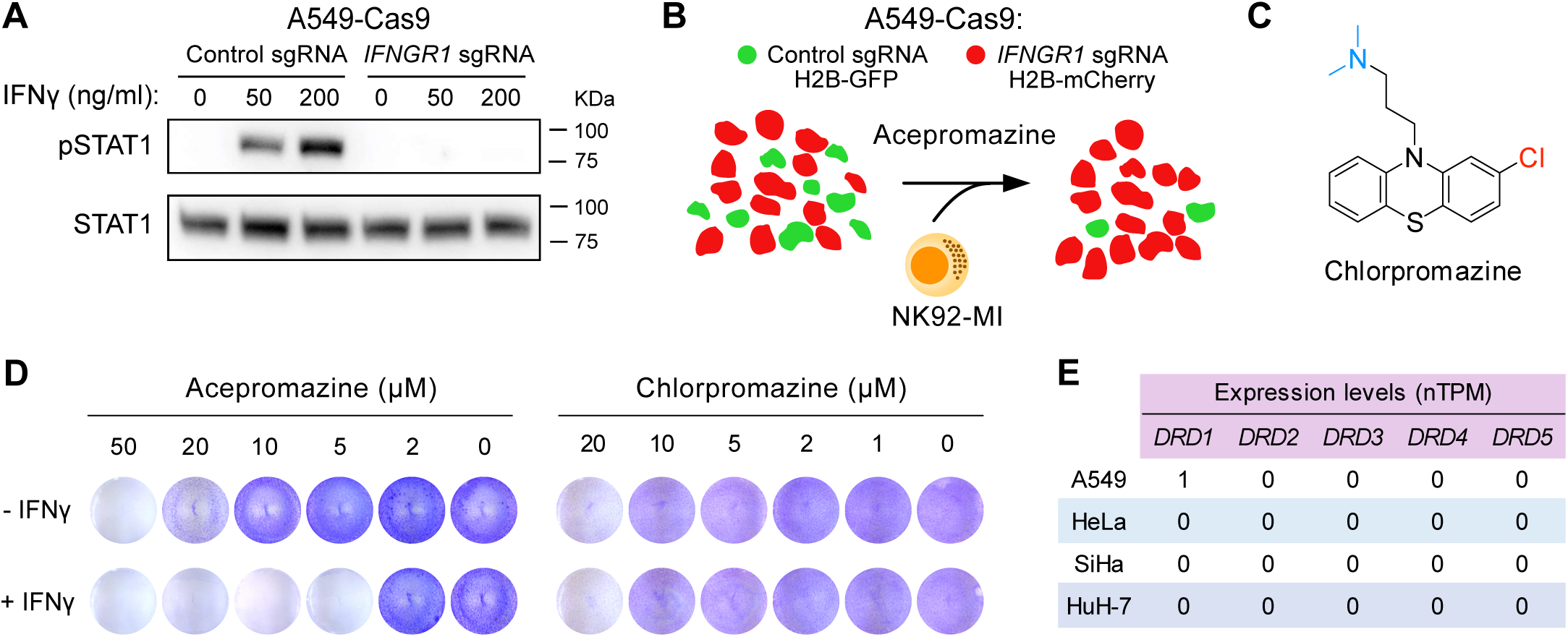
Acepromazine exhibits an interferon-enhanced selective anticancer activity independent of dopamine receptors. (A) Immunoblots of pSTAT1 and STAT1 in A549-Cas9 cells transduced with indicated sgRNAs and treated with IFNγ for 4 h. A representative result was shown from two independent experiments. Uncropped western blot images are provided as a Source Data file. (B) Schematic of the NK92-MI killing assay. An equal mixture of WT A549 cells (labeled with H2B-GFP) and *IFNGR1*-deficient A549 cells (labeled with H2B-mCherry) were co-cultured with NK-92MI cells for 48 h followed by FACS to quantify the ratio of GFP^+^ nuclei to mCherry^+^ nuclei. (C) The chemical structure of chlorpromazine. (D) Crystal violet staining of A549 (with or without 20 ng/ml IFNγ) treated with different concentrations of acepromazine or chlorpromazine. (E) Expression levels of genes encoding dopamine receptors in indicated acepromazine-sensitive cancer cell lines. Normalized transcript expression values, denoted nTPM, were obtained from the human protein atlas.

**Figure S2:**
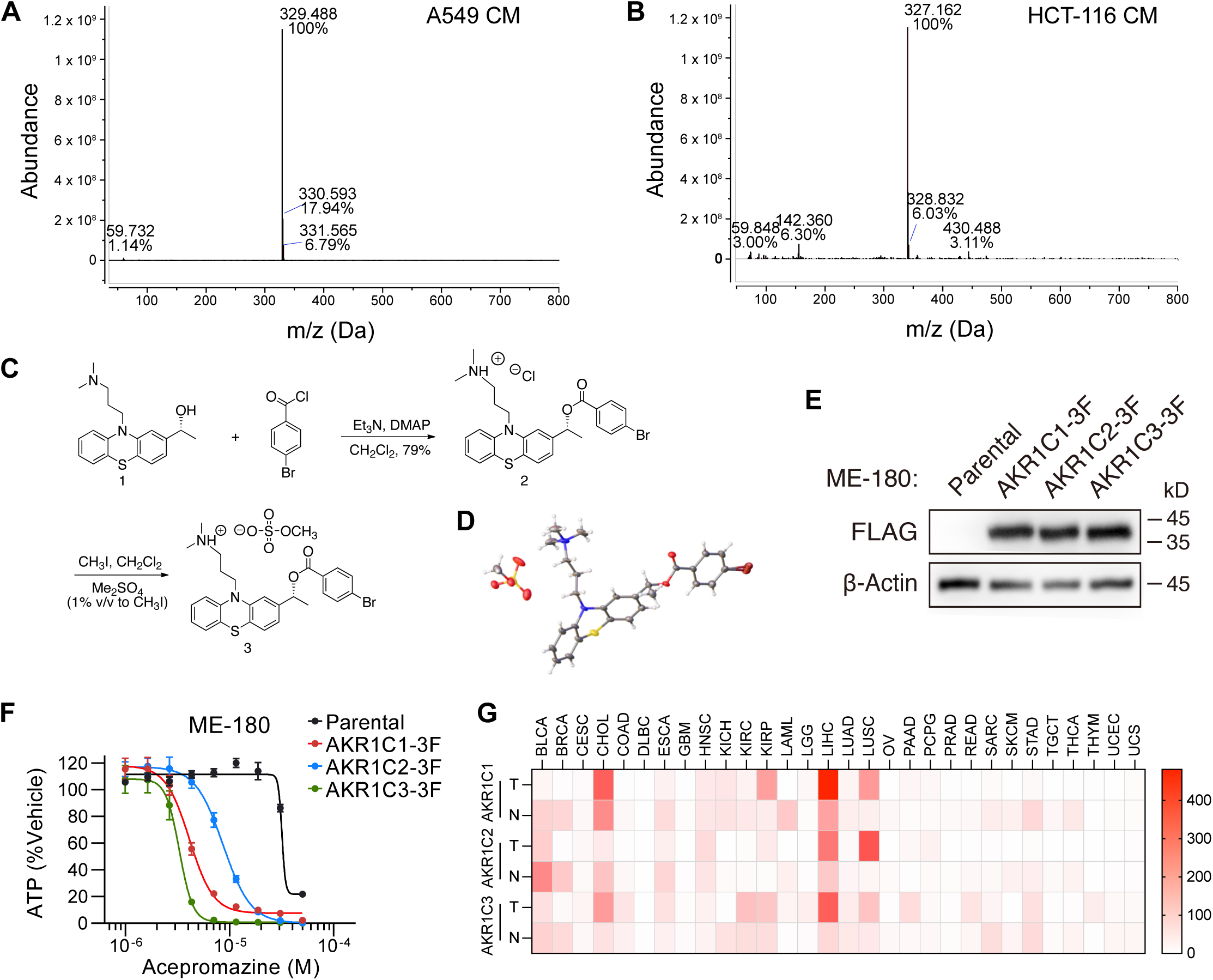
Metabolic activation of acepromazine into its stereo-selective active metabolite (*S*)-ACE-OH. (A) Mass spectrum of extracted metabolites from A549 conditioned medium. The dominant peak corresponds to the [M + H]^+^ of ACE-OH. (B) Mass spectrum of extracted metabolites from HCT-116 conditioned medium. The dominant peak corresponds to the [M + H]^+^ of ACE. (C) Synthesis of Compound 3 from the inactive enantiomer of ACE-OH (Compound 1). (D) Three-dimensional structure of Compound 3 determined by X-ray crystallography. Gray: carbon; white: hydrogen; blue: nitrogen; yellow: sulfur; red: oxygen; dark red: bromine. (E) Detection of ectopically expressed AKR1C1/2/3-3xFLAG in ME-180 cells by anti-FLAG western blotting. A representative result was shown from two independent experiments. Uncropped western blot images are provided as a Source Data file. (F) Concentration-response curves of ACE on the viability of ME-180 cells expressing AKR1C1/2/3-3xFLAG. Data represent the mean ± s.e.m. of three independent samples. IC_50_ and 95% CI are shown in Table S1. (G) Heatmap showing the median expression levels of genes encoding AKR1C1/2/3 in indicated tumor types and normal tissues. Median TPM (transcripts per million) values were downloaded from GEPIA 2.

**Figure S3:**
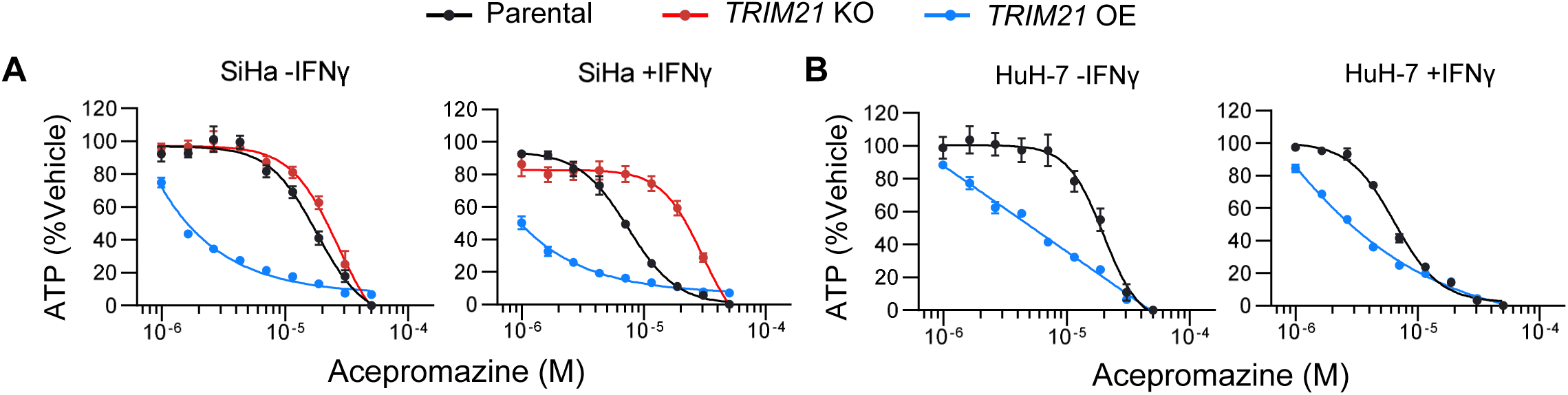
TRIM21 mediates the anticancer activity of acepromazine in multiple cancer cell lines. (A-B) Concentration-response curves of ACE on the viability of SiHa and HuH-7 cells with indicated genotypes with or without IFNγ (10 ng/ml) treatment. Data represent the mean ± s.e.m. of three independent samples. IC50 and 95% CI are shown in Table S1.

**Figure S4:**
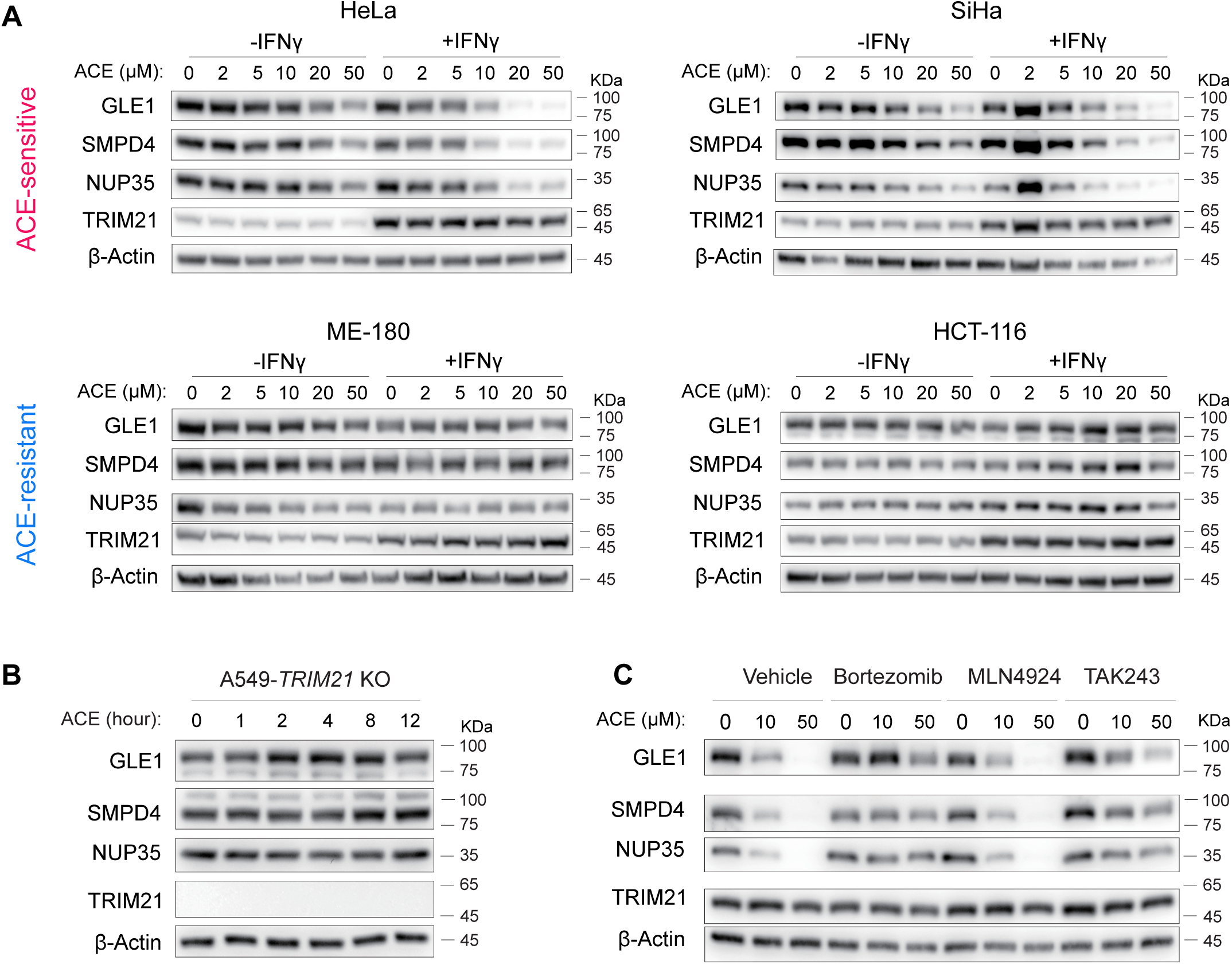
Acepromazine induces TRIM21-dependent degradation of nuclear pore proteins in multiple cancer cell lines. (A) Immunoblots of indicated proteins in ACE-sensitive or insensitive cell lines treated with indicated concentrations of ACE for 12 h with or without IFNγ (10 ng/ml) pretreatment. A representative result was shown from three independent experiments. Uncropped western blot images are provided as a Source Data file. (B) Immunoblots of indicated proteins in A549-*TRIM21* KO cells treated with ACE (20 μM) for indicated time. Uncropped western blot images are provided as a Source Data file. (C) Immunoblots of indicated proteins in IFNγ-stimulated A549 cells that were pretreated with bortezomib (100 nM), MLN4924 (200 nM), or TAK243 (200 nM) for 2 h followed by ACE (10 µM) treatment for 12 h. A representative result was shown from three independent experiments. Uncropped western blot images are provided as a Source Data file.

**Figure S5:**
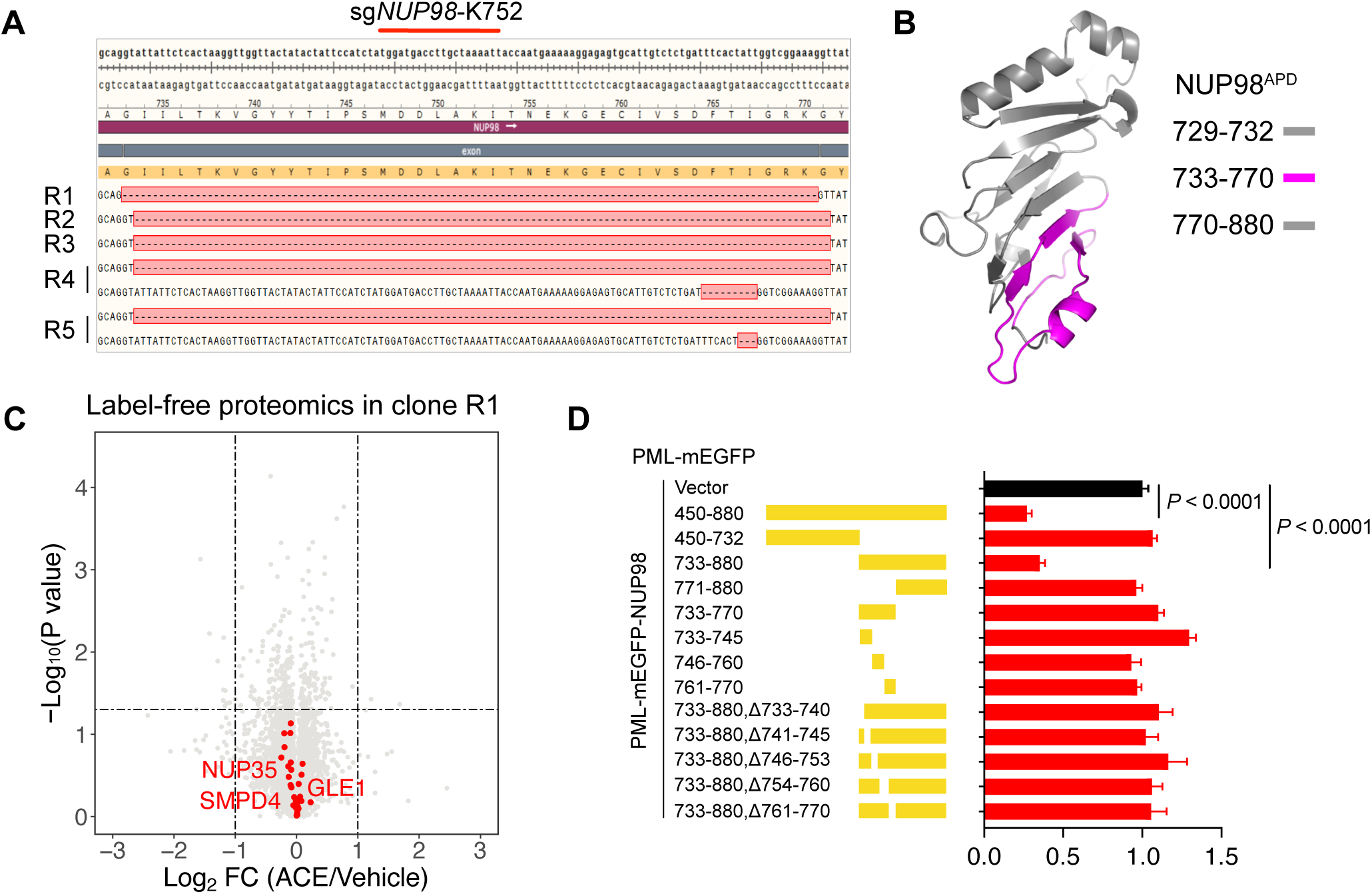
Deletions in NUP98^APD^ render acepromazine resistance. (A) The cDNA sequences of *NUP98* mutant clones. (B) The deleted segment in *NUP98* mutant clones highlighted in the structure of NUP98^APD^ (PDB: 2Q5Y). (C) Volcano plot depicting log_2_ transformed average fold change of each protein (quantified by label-free proteomics) and −log_10_ transformed *P* value comparing the *NUP98* mutant clone (R1) treated with ACE (10 µM) versus vehicle for 8 h. Three independent samples were included in the analyses. *P* values were calculated by unpaired Student’s t-test (two tailed). Data used for the plots are provided in Table S8. (D) The effect of ACE on the levels of indicated PML-GFP-NUP98 fusion proteins in A549-TRIM21^D355A^ cells. Data are the mean ± s.e.m. of six independent samples.

**Figure S6:**
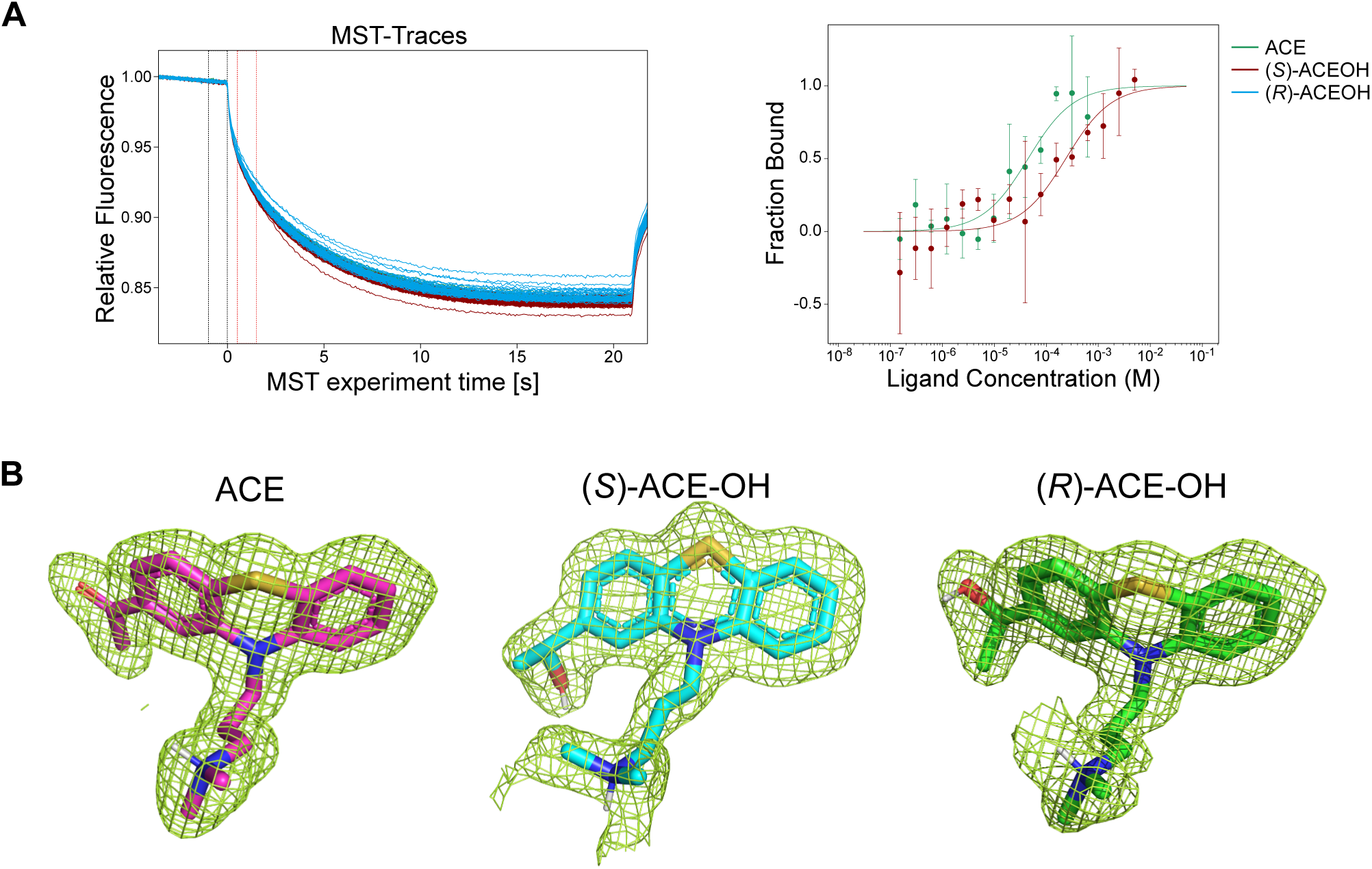
Characterization of the interaction of ACE and its metabolites with TRIM21 PRYSPRY. (A) The binding of ACE and its metabolites to the PRYSPRY domain of TRIM21^WT^ measured by MST. Raw MST traces and curve fitting are shown. (B) Electron density maps showing details of the regions occupied by indicated ligands in the PRYSPRY domain of TRIM21^D355A^. *Fo*-*Fc* omit map of ACE and its metabolites contoured at 3σ.

**Figure S7:**
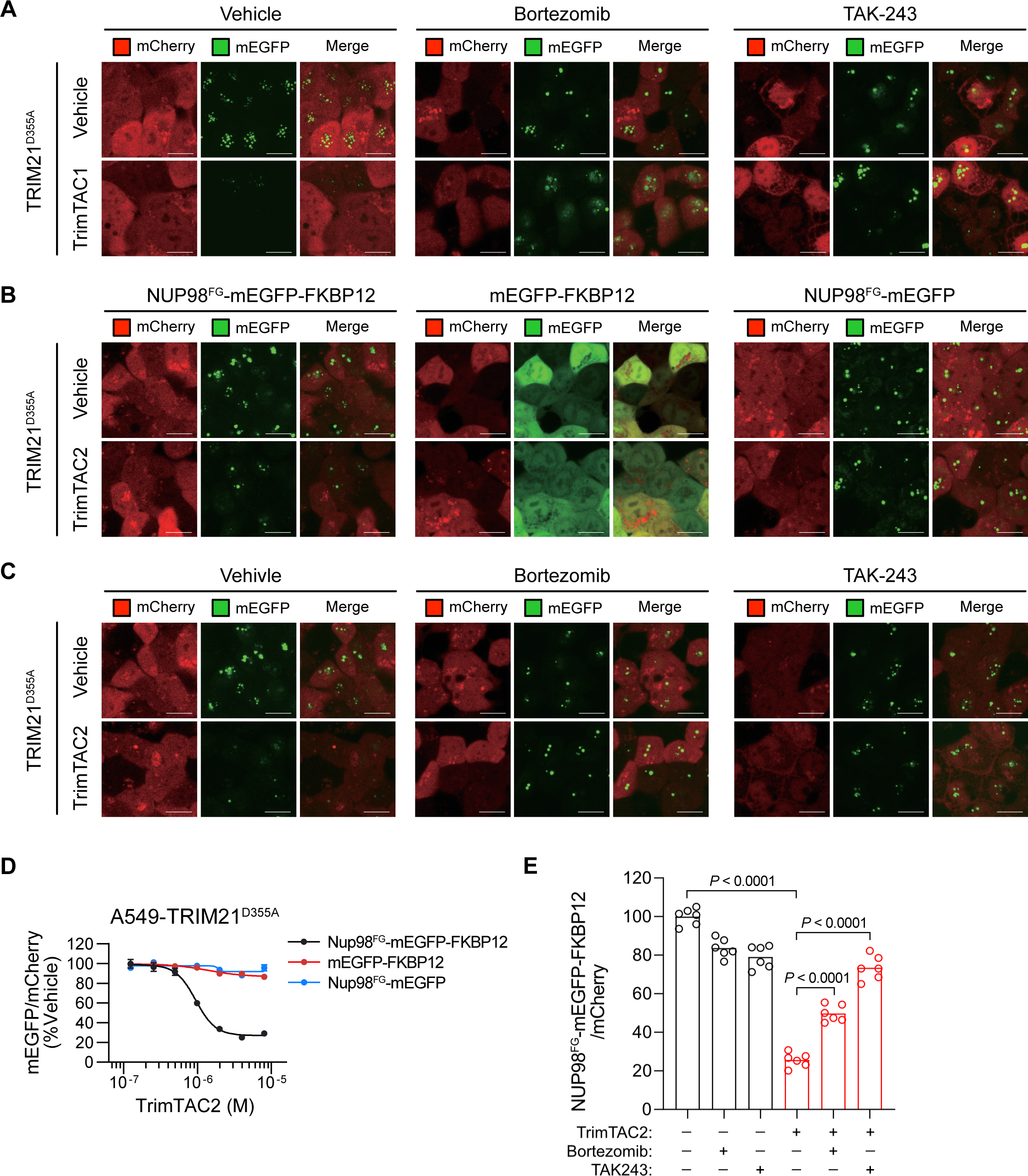
Design and characterization of TrimTACs. (A) Representative images of A549 cells expressing NUP98^FG^-mEGFP-BRD4^BD2^ pretreated with vehicle, bortezomib (400 nM), or TAK-243 (1.5 µM) for 2 h followed by TrimTAC1 (4 µM) treatment for 4 h. Scale bar: (B) Representative images of A549 cells expressing indicated reporter proteins treated with or without TrimTAC2 (10 µM) for 4 h. Scale bar: 10 μm. (C) Representative images of A549 cells expressing NUP98^FG^-mEGFP-FKBP12 pretreated with vehicle, bortezomib (200 nM), or TAK-243 (1 µM) for 2 h followed by TrimTAC2 (10 µM) treatment for 4 h. Scale bar: (D) Concentration-response curves of TrimTAC2 on the intensities of indicated reporter proteins in A549-TRIM21^D355A^ cells. Data indicate the mean ± s.e.m. of three independent samples. IC50 and 95% CI are shown in Table S1. (E) Intensities of NUP98^FG^-mEGFP-FKBP12 in A549-TRIM21^D355A^ cells with indicated treatment of TrimTAC2 (10 µM), bortezomib (200 nM), and TAK243 (1 µM). Data indicate the mean ± s.e.m. of six independent samples. One way ANOVA followed by Tukey’s multiple comparison test was used to determine statistical significance.

## Notes

### Competing Interest Statement

The authors have declared no competing interest.

## Main References

1 Stanton, B. Z., Chory, E. J. & Crabtree, G. R. Chemically induced proximity in biology and medicine. Science 359 (2018). 10.1126/science.aao5902

2 Wu, T. et al. Targeted protein degradation as a powerful research tool in basic biology and drug target discovery. Nat Struct Mol Biol 27, 605–614 (2020). 10.1038/s41594-020-0438-0

3 Bekes, M., Langley, D. R. & Crews, C. M. PROTAC targeted protein degraders: the past is prologue. Nat Rev Drug Discov 21, 181–200 (2022). 10.1038/s41573-021-00371-6

4 Krönke, J. et al. Lenalidomide causes selective degradation of IKZF1 and IKZF3 in multiple myeloma cells. Science 343, 301–305 (2014).

5 Lu, G. et al. The myeloma drug lenalidomide promotes the cereblon-dependent destruction of Ikaros proteins. Science 343, 305–309 (2014).

6 Liu, Y. et al. Expanding PROTACtable genome universe of E3 ligases. Nature Communications 14, 6509 (2023).

7 Chang, L., Ruiz, P., Ito, T. & Sellers, W. R. Targeting pan-essential genes in cancer: challenges and opportunities. Cancer cell 39, 466–479 (2021).

8 Zhang, X., Crowley, V. M., Wucherpfennig, T. G., Dix, M. M. & Cravatt, B. F. Electrophilic PROTACs that degrade nuclear proteins by engaging DCAF16. Nat Chem Biol 15, 737–746 (2019). 10.1038/s41589-019-0279-5

9 Guenette, R. G., Yang, S. W., Min, J., Pei, B. & Potts, P. R. Target and tissue selectivity of PROTAC degraders. Chem Soc Rev 51, 5740–5756 (2022). 10.1039/d2cs00200k

10 Hoegenauer, K. et al. Discovery of Ligands for TRIM58, a Novel Tissue-Selective E3 Ligase. ACS Medicinal Chemistry Letters 14, 1631–1639 (2023).

11 Buetow, L. & Huang, D. T. Structural insights into the catalysis and regulation of E3 ubiquitin ligases. Nature reviews Molecular cell biology 17, 626–642 (2016).

12 Alberti, S. & Hyman, A. A. Biomolecular condensates at the nexus of cellular stress, protein aggregation disease and ageing. Nat Rev Mol Cell Biol 22, 196–213 (2021). 10.1038/s41580-020-00326-6

13 Mitrea, D. M., Mittasch, M., Gomes, B. F., Klein, I. A. & Murcko, M. A. Modulating biomolecular condensates: a novel approach to drug discovery. Nat Rev Drug Discov 21, 841–862 (2022). 10.1038/s41573-022-00505-4

14 Shi, Y. et al. BRD4-targeting PROTAC as a unique tool to study biomolecular condensates. Cell Discovery 9, 1–13 (2023).

15 Castro, F., Cardoso, A. P., Gonçalves, R. M., Serre, K. & Oliveira, M. J. Interferon-gamma at the crossroads of tumor immune surveillance or evasion. Frontiers in immunology 9, 847 (2018).

16 Klingemann, H. The NK-92 cell line—30 years later: its impact on natural killer cell research and treatment of cancer. Cytotherapy (2023).

17 Collard, J. F. & Maggs, R. Clinical trial of acepromazine maleate in chronic schizophrenia. British Medical Journal 1, 1452 (1958).

18 Seibert, L. & Crowell-Davis, S. Antipsychotics. Veterinary Psychopharmacology, 201–215 (2019).

19 Ban, T. A. Fifty years chlorpromazine: a historical perspective. Neuropsychiatric disease and treatment 3, 495–500 (2007).

20 Wieder, M. et al. Identification of acepromazine and its metabolites in horse plasma and urine by LC–MS/MS and accurate mass measurement. Chromatographia 75, 635–643 (2012).

21 Barski, O. A., Tipparaju, S. M. & Bhatnagar, A. The aldo-keto reductase superfamily and its role in drug metabolism and detoxification. Drug metabolism reviews 40, 553–624 (2008).

22 Doench, J. G. et al. Optimized sgRNA design to maximize activity and minimize off-target effects of CRISPR-Cas9. Nature biotechnology 34, 184–191 (2016).

23 Li, W. et al. MAGeCK enables robust identification of essential genes from genome-scale CRISPR/Cas9 knockout screens. Genome biology 15, 1–12 (2014).

24 Foss, S. et al. TRIM21—from intracellular immunity to therapy. Frontiers in Immunology 10, 2049 (2019).

25 Ozato, K., Shin, D.-M., Chang, T.-H. & Morse III, H. C. TRIM family proteins and their emerging roles in innate immunity. Nature reviews immunology 8, 849–860 (2008).

26 Mallery, D. L. et al. Antibodies mediate intracellular immunity through tripartite motif-containing 21 (TRIM21). Proc Natl Acad Sci U S A 107, 19985–19990 (2010). 10.1073/pnas.1014074107

27 McEwan, W. A. et al. Intracellular antibody-bound pathogens stimulate immune signaling via the Fc receptor TRIM21. Nat Immunol 14, 327–336 (2013). 10.1038/ni.2548

28 James, L. C., Keeble, A. H., Khan, Z., Rhodes, D. A. & Trowsdale, J. Structural basis for PRYSPRY-mediated tripartite motif (TRIM) protein function. Proc Natl Acad Sci U S A 104, 6200–6205 (2007). 10.1073/pnas.0609174104

29 Hampoelz, B., Andres-Pons, A., Kastritis, P. & Beck, M. Structure and assembly of the nuclear pore complex. Annual review of biophysics 48, 515–536 (2019).

30 Branon, T. C. et al. Efficient proximity labeling in living cells and organisms with TurboID. Nature biotechnology 36, 880–887 (2018).

31 Vinyard, M. E. et al. CRISPR-suppressor scanning reveals a nonenzymatic role of LSD1 in AML. Nature chemical biology 15, 529–539 (2019).

32 Gosavi, P. M. et al. Profiling the landscape of drug resistance mutations in neosubstrates to molecular glue degraders. ACS Central Science 8, 417–429 (2022).

33 Rosenblum, J. S. & Blobel, G. Autoproteolysis in nucleoporin biogenesis. Proceedings of the National Academy of Sciences 96, 11370–11375 (1999).

34 Lallemand-Breitenbach, V. & de Thé, H. PML nuclear bodies: from architecture to function. Current opinion in cell biology 52, 154–161 (2018).

35 Winter, G. E. et al. Phthalimide conjugation as a strategy for in vivo target protein degradation. Science 348, 1376–1381 (2015).

36 Clift, D. et al. A Method for the Acute and Rapid Degradation of Endogenous Proteins. Cell 171, 1692–1706 e1618 (2017). 10.1016/j.cell.2017.10.033

37 Zeng, J. et al. Target-induced clustering activates Trim-Away of pathogens and proteins. Nat Struct Mol Biol 28, 278–289 (2021). 10.1038/s41594-021-00560-2

38 Amara, J. F. et al. A versatile synthetic dimerizer for the regulation of protein–protein interactions. Proceedings of the national academy of sciences 94, 10618–10623 (1997).

39 Banani, S. F. et al. Compositional control of phase-separated cellular bodies. Cell 166, 651–663 (2016).

40 Courchaine, E. et al. The coilin N-terminus mediates multivalent interactions between coilin and Nopp140 to form and maintain Cajal bodies. Nature Communications 13, 6005 (2022).

41 Mitrea, D. M. et al. Self-interaction of NPM1 modulates multiple mechanisms of liquid– liquid phase separation. Nature communications 9, 842 (2018).

42 Ross, C. A. & Poirier, M. A. Protein aggregation and neurodegenerative disease. Nature medicine 10, S10–S17 (2004).

43 Taniue, K. & Akimitsu, N. Aberrant phase separation and cancer. The FEBS journal 289, 17–39 (2022).

44 Xia, S., Chen, Z., Shen, C. & Fu, T.-M. Higher-order assemblies in immune signaling: supramolecular complexes and phase separation. Protein & cell 12, 680–694 (2021).

45 D’Amico, F., Mukhopadhyay, R., Ovaa, H. & Mulder, M. P. Targeting TRIM proteins: a quest towards drugging an emerging protein class. ChemBioChem 22, 2011–2031 (2021).

## Methods References

46 Lv, L. et al. Discovery of a molecular glue promoting CDK12-DDB1 interaction to trigger cyclin K degradation. Elife 9 (2020). 10.7554/eLife.59994

47 Hughes, C. S. et al. Single-pot, solid-phase-enhanced sample preparation for proteomics experiments. Nature protocols 14, 68–85 (2019).

48 Hou, J., Wang, J., Yang, F. & Xu, T. DIA-MS2pep: a library-free framework for comprehensive peptide identification from data-independent acquisition data. Biophysics Reports 8, 253 (2022).

49 Spivak, M., Weston, J., Bottou, L., Kall, L. & Noble, W. S. Improvements to the percolator algorithm for peptide identification from shotgun proteomics data sets. Journal of proteome research 8, 3737–3745 (2009).

50 Demichev, V., Messner, C. B., Vernardis, S. I., Lilley, K. S. & Ralser, M. DIA-NN: neural networks and interference correction enable deep proteome coverage in high throughput. Nature methods 17, 41–44 (2020).

51 Joung, J. et al. Genome-scale CRISPR-Cas9 knockout and transcriptional activation screening. Nat Protoc 12, 828–863 (2017). 10.1038/nprot.2017.016

52 Wang, Q.-S. et al. The macromolecular crystallography beamline of SSRF. Nucl. Sci. Tech 26, 12–17 (2015).

53 Wang, Q.-S. et al. Upgrade of macromolecular crystallography beamline BL17U1 at SSRF. Nuclear Science and Techniques 29, 1–7 (2018).

54 Kabsch, W. xds. Acta Crystallographica Section D: Biological Crystallography 66, 125–132 (2010).

55 Evans, P. R. An introduction to data reduction: space-group determination, scaling and intensity statistics. Acta Crystallographica Section D: Biological Crystallography 67, 282–292 (2011).

56 Evans, P. R. & Murshudov, G. N. How good are my data and what is the resolution? Acta Crystallographica Section D: Biological Crystallography 69, 1204–1214 (2013).

57 McCoy, A. J. et al. Phaser crystallographic software. Journal of applied crystallography 40, 658–674 (2007).

58 Emsley, P., Lohkamp, B., Scott, W. G. & Cowtan, K. Features and development of Coot. Acta Crystallographica Section D: Biological Crystallography 66, 486–501 (2010).

59 Afonine, P. V. et al. Towards automated crystallographic structure refinement with phenix. refine. Acta Crystallographica Section D: Biological Crystallography 68, 352–367 (2012)

